# Solution structure and synaptic analyses reveal determinants of bispecific T cell engager potency

**DOI:** 10.1101/2022.06.15.496334

**Authors:** Alexander Leithner, Oskar Staufer, Tanmay Mitra, Falk Liberta, Salvatore Valvo, Mikhail Kutuzov, Hannah Dada, Jacob Spaeth, Sally Zhou, Felix Schiele, Sophia Reindl, Herbert Nar, Stefan Hoerer, Maureen Crames, Stephen Comeau, David Young, Sarah Low, Edward Jenkins, Simon J. Davis, David Klenerman, Andrew Nixon, Noah Pefaur, David Wyatt, Omer Dushek, Srinath Kasturirangan, Michael L. Dustin

**Affiliations:** The Kennedy Institute of Rheumatology, University of Oxford, Oxford, OX3 7FY, United Kingdom; Chinese Academy of Medical Science Oxford Institute, University of Oxford, OX3 7BN, United Kingdom; Boehringer Ingelheim Pharma GmbH & CO. KG, Birkendorfer Strasse 65, 88397 Biberach an der Riss, Germany; Sir William Dunn School of Pathology, University of Oxford, Oxford, OX1 3RE, United Kingdom; Biotherapeutics Discovery, Boehringer Ingelheim, Ridgefield, CT, 06877, USA; Medical Research Council Human Immunology Unit, University of Oxford, Oxford, OX3 9DU, United Kingdom; Radcliffe Department of Medicine, University of Oxford, Oxford, OX3 9DU, United Kingdom; Yusuf Hamied Department of Chemistry, University of Cambridge, Cambridge, CB2 1EW, United Kingdom

**Author notes:** contributed equally (Alexander Leithner, Oskar Staufer). contributed equally (Tanmay Mitra, Falk Liberta). Helmholtz Institute for Pharmaceutical Research Saarland, Saarbruecken, Germany (Oskar Staufer).

**Keywords:** Immunological Synapse, Breast Cancer, Supported Lipid Bilayers, Immunotherapy, Multi-state Modelling

## Abstract

Bispecific T-cell engagers (TcEs) link T cell receptors to tumor-associated antigens on cancer cells, forming cytotoxic immunological synapses (IS). Close membrane-to-membrane contact (≤13 nm) has been proposed as a key mechanism of TcE function. To investigate this and identify potential additional mechanisms, we compared four immunoglobulin G1-based (IgG1) TcE Formats (A-D) targeting CD3ε and Her2, designed to create varying intermembrane distances (A<B<C<D). Small-angle X-ray scattering (SAXS) and modelling of the conformational states of isolated TcEs and TcE-antigen complexes predicted close-contacts (≤13 nm) for Formats A and B and far-contacts (≥18 nm) for Formats C and D. In supported lipid bilayer (SLB) model interfaces, Formats A and B recruited, whereas Formats C and D repelled, CD2-CD58 interactions. Formats A and B also excluded bulky Quantum dots more effectively. SAXS also revealed that TcE-antigen complexes formed by Formats A and C were less flexible than complexes formed by Formats B and D. Functional data with Her2-expressing tumor cells showed cytotoxicity, surface marker expression and cytokine release following the order A>B=C>D. In a minimal system for IS formation on SLBs, TcE performance followed the trend A=B=C>D. Addition of close-contact requiring CD58 co-stimulation revealed phospholipase C-γ activation matching cytotoxicity with A>B=C>D. Our findings suggest that, when adhesion is equivalent, TcE potency is determined by two parameters: contact distance and flexibility. Both the close/far-contact formation axis and the low/high flexibility axis significantly impact TcE potency, explaining the similar potency of Format B (close-contact/high flexibility) and C (far-contact/low flexibility).

**Significance statement:** Bispecific T-cell engagers (TcEs) are immunotherapeutic drugs that trigger the destruction of cancer cells by linking T cells to cancer cell through specific surface molecules (antigens). We designed a series of TcEs with varying distances between their binding sites and flexibilities of the TcE-antigen complexes. By combining structural and functional analyses, we confirmed close-contact formation between T cells and cancer cells as a critical determinant, mediated by co-activating receptors. Furthermore, we also identified molecular flexibility of the TcE-antigen complex as a further critical parameter for TcE potency. These findings provide novel insights into TcE function and highlight the importance of both parameters for future research and the design of improved immunotherapies.

## Introduction

Bispecific T cell engagers (TcEs) that are able to simultaneously bind to epitopes on cancer cells and T cells, are a versatile, highly-potent and economical immunotherapeutic tool (1). TcEs are antibody-derived molecules harboring at least one antigen binding site (paratope) that is selective for T cells, typically directed against CD3ε of the T cell receptor (TCR) complex, while the other paratope is targeting a tumor-associated antigen (TAA). A plethora of TcE-designs have been developed (2), which can be subcategorized into those that retain the fragment-crystallizable (Fc) region of immunoglobulins (IgGs) and those that only contain minimal antigen binding arms, influencing their pharmacokinetic properties and biodistribution. In both cases, the paratope carrying units are typically linked by at least one flexible hinge or spacer.

Mechanistically, TcEs physically link polyclonal effector T cells with cancer cells to promote the formation of cytotoxic immunological synapses (ISs), exploiting the intrinsic cytotoxic machinery of T cells for a therapeutic benefit (3). Since the first-in-class approval of blinatumomab in 2015, a large variety of different TcE formats were (pre-) clinically assessed in solid and hematological tumors (4), particularly for lymphoblastic leukemia and Her2^+^ solid tumors (5). These assessments revealed significant variations in TcE safety and efficacy, which is determined by a variety of factors such as the abundance and accessibility of TAA on target cells (6, 7) as well as TcE-intrinsic parameters such as the orientation and spacing of paratopes as well as their binding affinity, valency and Fc-design (4, 8).

IS formation between CD8^+^ T cells and target cells is a highly orchestrated process, dominated by a complex interplay of molecular self-assembly events and active cellular rearrangements. IS structure, maturation and signaling have been reviewed in detail elsewhere (9). Central structural elements underlying antigen recognition and signal integration at the IS include TCR and filamentous-actin (F-actin)-enriched microclusters and a central supramolecular activation cluster (cSMAC) adjacent to an F-actin depleted secretory domain, integrin lymphocyte function-associated antigen 1 (LFA-1) – intercellular adhesion molecule 1 (ICAM1) based peripheral SMAC (pSMAC) and a distal (dSMAC) region that is a site of F-actin polymerization (10). Additionally, the dSMAC also accumulates interactions of the co-stimulatory receptor CD2 on T cells and its ligand CD58, forming a corolla of microdomains (11) separated by a bulk phase enriched in glycoproteins with large ectodomains (ECDs), such as CD45 and CD43. Importantly, IS formation can be modelled using supported lipid bilayers (SLBs) that present peptide-major histocompatibility complex (pMHC) or anti-CD3 antibodies, along with ICAM1 and CD58, which serve as ligands for adhesion molecules. This well-established approach effectively recapitulates SMAC formation and allows for the analysis of signaling processes with a resolution and sensitivity that remains unmatched in intercellular synapses (12).

TCR intrinsic and extrinsic mechanisms are likely to cooperate in T cell activation across a range of conditions (13). Intrinsic mechanisms include clustering and conformational change. TCR clustering by multivalent soluble ligands activates signaling, a process that is notably required by chimeric antigen receptors (14). Conformational changes are readily demonstrated in the cytoplasmic domains of the TCR and allosteric coupling of the extracellular domains to the transmembrane domains has been demonstrated (15, 16). Catch-bonds represent another manifestation of TCR-pMHC conformational changes, where forces on the order of 10 pN lead to a decrease in the dissociation rate (k_off_), thereby creating greater opportunity for multistep signaling processes governed by kinetic proofreading (17, 18). Extrinsic mechanisms include the role of Csk, a negative regulator of the key Src family kinase Lck, where inhibition of Csk leads to TCR signaling (19). Additionally, the exclusion of the large inhibitory transmembrane phosphatase CD45 from close-contacts (∼13 nm intermembrane distance) established by the interaction of TCR-pMHC and co-stimulatory receptor-ligand pairs, such as CD2-CD58, promotes TCR signaling as summarized in the kinetic segregation (KS) model (20, 21). Notably, the interplay of these alternative mechanisms may contribute to the robustness of TCR triggering. For example, sufficient TCR clustering can compensate for the absence of CD45 exclusion (22).

In the context of TcEs, multiple studies have found that targeting membrane-proximal epitopes (23–25) leads to the highest potency and is accompanied by CD45 exclusion (26). Notably, comparing a full-length IgG2-based TcE with smaller formats revealed that smaller formats are more efficient against distal epitopes but face steric barriers when trying to access membrane proximal epitopes (27), suggesting an optimal close-contact for TcE performance. These controversies highlight that several questions of fundamental relevance for TcE-based therapeutic design remain: to what degree is the canonical IS organization maintained in the TcE-mediated IS? How does TcE design impact on the dynamics of IS formation, TCR-signaling strength and the formation of tight membrane-to-membrane interactions? How do different TcE formats synergize with co-stimulatory receptors and arrange them in space? And: are there additional parameters beyond close-contacts and clustering that govern TcE performance?

To address these questions, we combined structural analysis of CD3ε-TcE-TAA complexes in solution through small-angle X-ray scattering (SAXS), functional cell-based assays, and a reductionistic SLB system that we adapted to present Her2 as a model TAA. This approach enabled us to study the contacts formed between T cells or T cell-mimicking giant unilamellar vesicles (GUVs) and cancer cell-mimicking SLBs. We focused on a set of four well-characterized modular TcEs with varying paratope spacing and positioning, allowing us to extract the quantitative parameters underpinning signaling at the TcE-mediated IS. We found that in-solution antigen spacing and flexibility and the analysis of TcE-mediated membrane apposition provide complementary information. We confirm close-contact formation as an important determinant for TcE performance and uncover the direct recruitment of CD2-CD58 co-stimulation to the site of TcE intermembrane assemblies as one mechanism underlying its potency. Furthermore, we identify molecular flexibility of TcE-antigen complexes as a new parameter of interest to determine TcE potency. Our data suggest that the effects of flexibility and close-contact formation are on similar orders of magnitude.

## Results

We began our study by designing a panel of four distinct bispecific TcEs, termed Formats A-D, by using immunoglobulin modules to increase the span between the paratopes of the TcEs in ∼ 4 nm steps (**Fig. 1A**). The TcEs were engineered using knob-into-hole (28) technology, incorporating single-chain variable (scFv) and single-chain antigen-binding fragments (scFab) to enable monovalent binding to the membrane proximal domain IV of human Her2 (29) on tumor cells, based on trastuzumab, and the C-terminal peptide (30) of human CD3ε on T cells. All TcEs were confirmed to be monomeric by size exclusion chromatography with the expected hydrodynamic radii **(Supplementary Figure 1A and F).** TcEs that included scFvs - namely, A, B and D, exhibited lower thermal stability based on differential scanning calorimetric analysis and greater hydrophobicity as indicated by hydrophobic interaction chromatography **(Supplementary Figure 1B and C).** Surface plasmon resonance indicated that all TcEs had an affinity for CD3ε in the low nanomolar range (<10 nM), similar to previously reported affinities for CD3εδ (31) and CD3εγ heterodimers. TcE affinities for Her2 ranged from ∼80 to 250 pM, consistent with published values for trastuzumab (32, 33) **(Supplementary Figure 1D-F)**.

**Figure 1.**
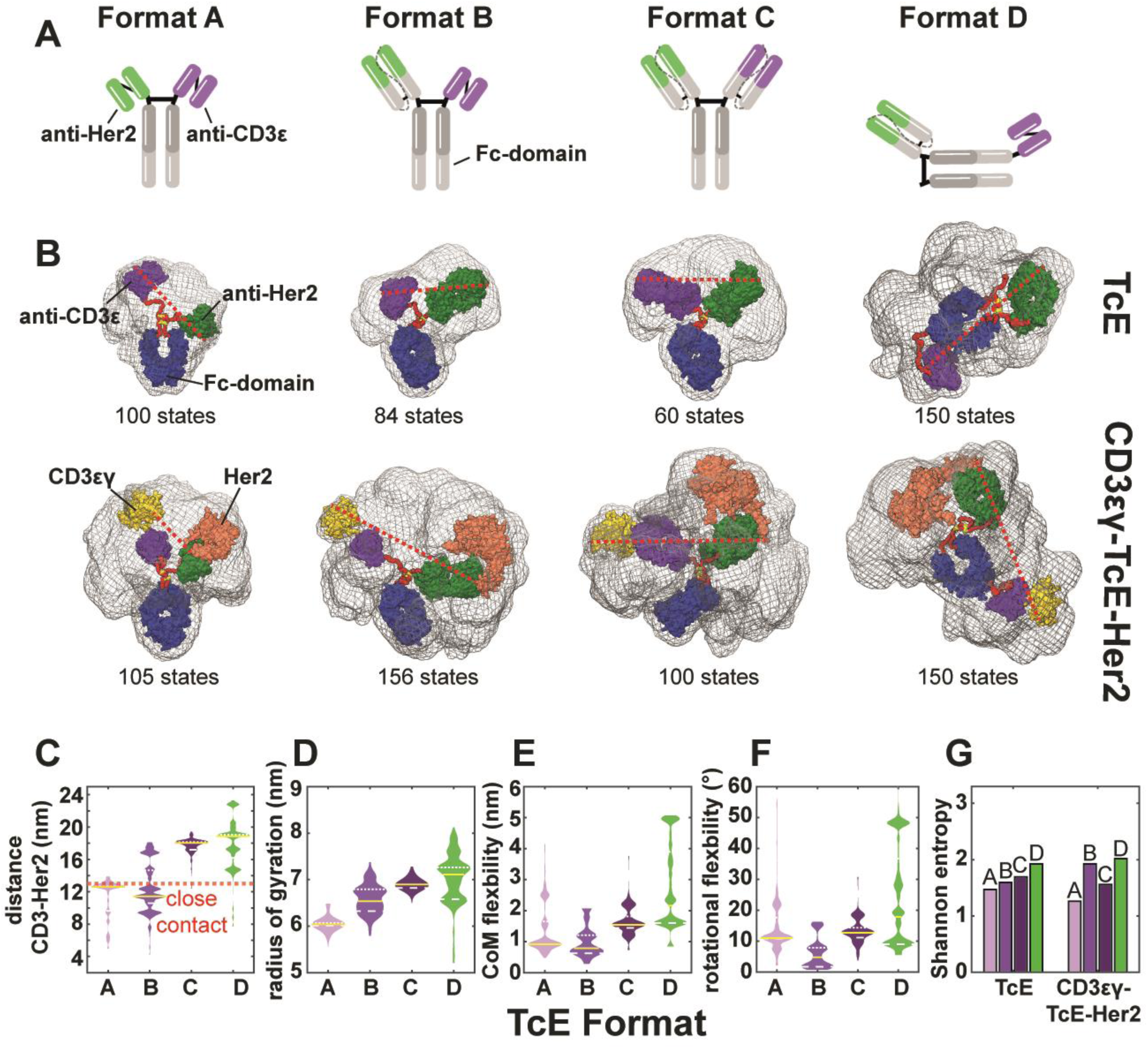
Structural analysis of TcEs. (**A**) Schematic overview of TcE formats. (**B**) Conformational space of TcEs (top) and TcE-antigen complexes (bottom) in solution. Grey meshes show overlay of all conformational states of each molecule. One single state per molecule is shown as colored surface representation. Magenta = anti-CD3, green = anti-Her2, blue = F_c_-domain, yellow = CD3εγ, salmon = Her2. Flexible residues are shown in red. Disulphide bonds in the hinge region are shown in yellow. Dashed red line illustrates measurement of the distance between antigens. (**C**) SAXS-data derived conformation weight-adjusted distributions of antigen spacings, (**D**) radius of gyration, (**E**) center of mass (CoM) flexibility, (**F**) rotational flexibility and (**G**) Shannon entropy. Solid yellow lines indicate medians, white dotted lines represent 1^st^ and 3^rd^ quartiles respectively, red dashed lines indicate close contact zone at 13 nm.

While we designed the TcE-series to increase paratope spacing in 4 nm increments, the actual range of spacing between paratopes and between the antigens in TcE-antigen complexes is difficult to predict. Thus, we decided to utilize small-angle X-ray scattering (SAXS) to assess size and intramolecular spacing in our TcE-series (34). SAXS data were either collected from TcEs alone or from heterotrimers formed between TcEs and the ectodomains of Her2 and CD3εγ. The collected data (**Supplementary Fig. 1G)** were analyzed by multi-state modelling, resulting in 60-156 states per TcE or heterotrimer, representing the conformational space that is covered by the respective TcE or TcE-antigen complex in solution (**Fig 1B and Supplementary Fig. 1H)**. This was followed by measurements for every single state, either of the direct distances between the antibody-heavy chain complementarity-determining region 3 (CDR3) loops or of the distances between the C-terminal sites that would anchor CD3ε and Her2 to the respective cellular membranes, to obtain paratope- or anchor-to-anchor spacing profiles, respectively. Importantly, paratope spacing did not follow a simple incremental increase from Format A to D. Each individual TcE alone produced a non-normal distribution of paratope-spacings between 3 and 16 nm, with Formats A and B having similar values of 8.8 and 9.6 nm respectively, and Formats C and D having a similar median spacing of 14.6 and 14.5 nm, respectively (**Supplementary Fig. 2A)**. Modelling of the CD3ε-TcE-Her2 complexes showed that Formats A and B converged around similar anchor-to-anchor distances of 12.6 and 11.4 nm, while Formats C and D had median anchor-to-anchor distances of 18.0 and 18.9 nm, respectively (**Fig. 1C**). Notably, Formats A and B generated distances that are similar to the close-contact distances of 12.8 and 13.1 nm measured for CD2-CD58 and TCR-pMHC interactions, respectively (20, 35). In contrast, Formats C and D exceed the close-contact standard by ∼ 5 nm. Notably, this results in significant functional impairment in experiments where the length of the TCR-pMHC complex is elongated (20), generating what we define here as far-contacts.

Multistate modelling also provides information about molecular flexibility (34), which has not been extensively investigated for TcEs. We analyzed measures of flexibility including radius of gyration, flexibility of center of mass (CoM Flex) and rotational flexibility, weighed by state-specific contribution across the multi-state ensembles. To quantify effective disorder, Shannon entropy, a measure of uncertainty within a distribution, was calculated by multiplying the weighted CoM Flex and rotational flexibilities (**Fig. 1D-G, Supplementary Figure 2B-F and Movies S1-4**). Interestingly, the binding of antigens to Format A resulted in a decrease in measures of flexibility suggesting conformational stabilization (**Fig. 1D-G and Supplementary Fig. 2B-E**). Notably, the consistently greater flexibility of Format D was also reflected in broader tilt angle distributions of intramolecular domains relative to the Her2-anti-Her2 axis (**Supplementary Fig. 2G**). Importantly, these findings were further corroborated by plotting the model-free SAXS data on dimensionless Kratky plots (36), which allow for the comparison of the shape and flexibility of proteins of different sizes, with greater flexibility indicated by a higher amplitude. Format D exhibited a higher amplitude, distinguishing it from the other formats, whether analyzed alone or in combination with antigens. Based on the Kratky plots, Format A became less flexible upon the addition of antigens, setting it apart from Formats B and C (**Supplementary Fig. 2H & I**). Importantly, relative flexibility assessed by the Kratky plots was similar to the calculated Shannon entropy from the multistate modelling for TcE-antigen complexes and TcEs in isolation (**Fig. 1G and Supplementary Fig. 2E)**. Taken together, our data suggest that our TcE-series can be further categorized based on flexibility with Format A forming less-flexible assemblies than B, within the close-contact-forming TcEs, and Format C forming less-flexible assemblies than D, within the far-contact-forming TcEs.

Since SAXS measurements are performed in solution, without the anchorage of the investigated proteins to the cell membrane, we aimed to corroborate our findings in a system that mimics the interaction of two biological membranes. To this end, we built a minimal system for membrane-to-membrane contacts, where a SLB was functionalized with fluorescently labeled and polyhistidine-tagged recombinant Her2 ectodomains *via* NTA(Ni^2+^) coupling. In addition, we added CD58, the ligand for co-stimulatory receptor CD2 on T cells, with a distinct fluorescent tag, to the SLB. Subsequently, we generated giant unilamellar vesicles (GUVs) (37) that we functionalized with unlabeled CD3εδ and CD2 protein, which were then seeded onto the SLB, preloaded with TcEs (**Fig. 2A**). CD2-CD58 spans a well-characterized distance of 12.8 nm (38) with limited flexibility, making it a *bona fide* molecular ruler (35). We hypothesized that the ability to colocalize interfacial CD2-CD58 interactions with the Her2-TcE-CD3εδ assemblies would depend on the TcEs’ anchor-to-anchor spacing predicted by SAXS. Supporting our hypothesis, we observed a mean positive correlation for Her2 and CD58 for Formats A and B, but mean negative correlations for C and D (**Fig. 2B & C)**. The negative correlation manifested as low Her2 recruitment, as shown, or low CD2-CD58 interaction within contacts enriched for Formats C and D. Notably, while the vast majority of GUVs showed homogenous accumulation and distribution of Her2 and CD58, we noticed that some GUVs exhibited segregated domains of the two proteins with multiple incidences for Formats C and D; with the highest frequency for Format D (**Supplementary Fig. 3A & B)**. This is in line with the SAXS data in which anchor-to-anchor spacing of Formats A and B is similar to the ∼13 nm gap that is formed by CD2-CD58, supporting close-contact and recruitment of CD2-CD58, whereas Formats C and D exceed this distance, supporting far-contact and repulsion of CD2-CD58.

**Figure 2.**
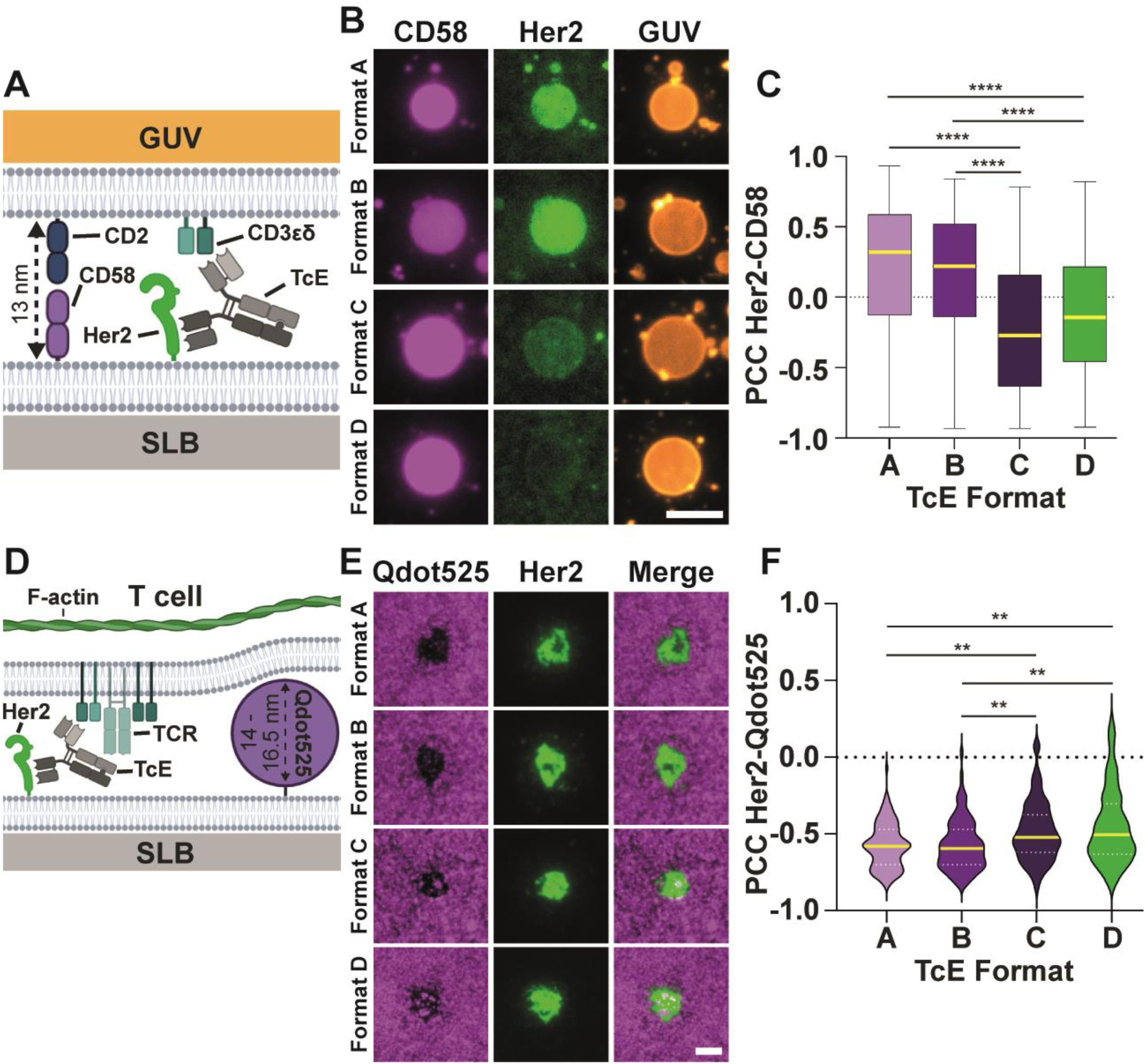
Intermembrane distances in membrane-model system. (**A**) Schematic overview of GUV-SLB experimental system. (**B**) Representative TIRF images of CD3εδ and CD2 loaded GUVs (orange) incubated for 60 min. on SLBs presenting 400 molecules/µm^2^ Her2 (green) and 200 molecules/µm^2^ CD58 (magenta), pre-incubated with TcEs, scale bar = 5µm. (**C**) Pearson correlation coefficient of Her2 and CD58 at the interface of CD3εδ and CD2 loaded GUVs and SLBs from (B). Yellow solid line = median, box and whiskers = min. and max. 3 biological replicates, Kruskal-Wallis test, **** p<0.0001. (**D**) Schematic overview of T cell-SLB experimental system in the presence of Qdot525. (**E**) Representative TIRF images of TcE-mediated contacts of T cells on SLBs presenting Her2 at 400 molecules/µm^2^ (green) and QDot525 (magenta), pre-incubated with TcEs, scale bar = 5µm. (**F**) Pearson correlation coefficient of Her2 and Qdot525 at TcE mediated T cell-SLB contacts from (E), yellow solid line = median, white dashed line = quartiles, 3 biological replicates, Kruskal-Wallis test, ** p≤0.0058.

To test the relevance of these findings in the context of T cells, but in the absence of any other adhesion system, we replaced CD58 with Quantum dots of a defined diameter of 14 – 16.5 nm (39, 40) (Qdot525) in the SLB (**Fig. 2D**) and measured their exclusion in regions of TcE-mediated Her2 clustering upon the addition of primary human T cells. While all formats induced considerable exclusion of Qdot525, we observed significant differences between the formats, with A=B>C=D (**Fig. 2E & F**). These results further validate that Formats A and B form close-contacts whereas Format C and D form far-contacts.

Taken together, the SAXS, GUV-SLB and T cell-SLB data demonstrated that our TcE-series can be categorized into two groups, based on the anchor-to-anchor spacing they mediate, with Formats A and B at ∼ 13 nm, and C and D at ∼ 18 nm. Within these pairs, B is more flexible than A, and D is more flexible than C. Thus, our TcE panel provides tools to study both contact distance and flexibility of the TcE-mediated molecular complex as critical parameters in TcE potency for a membrane-proximal epitope.

Next, we assessed how these parameters influence the cytotoxic potency of the different TcE formats in co-cultures of breast cancer cell-lines and primary, activated human CD8^+^ T cells. To evaluate the impact of Her2 surface expression on cancer cells, a crucial factor for TcE on-target/off-tumor targeting (41), we included two cell lines with differing Her2 membrane densities: BT474 ductal carcinoma breast cancer cells, with ∼6×10^5^ molecules Her2/cell (Her2^High^), and MCF-7 mammary epithelium adenocarcinoma cells, with ∼2×10^4^ Her2/cell (Her2^Low^) (42). We further verified that co-stimulatory ligands ICAM1, CD58 (LFA-3) and CD80 were expressed on both cell lines at comparable levels (**Supplementary Fig. 3C**). Cytotoxicity was measured using LDH release endpoint assays in an E:T ratio of 5:1 (T cell: cancer cell) over 24 hours. Both Her2^High^ and Her2^Low^ cells were killed effectively with all TcEs, but there was a trend for the effective median concentration (EC_50_), with A<B=C=D, as well as significant differences in the maximum killing levels, with A>B≥C>D (**Fig. 3A and B and Supplementary Fig. 3D & E**). Additionally, we performed time-lapse imaging of the first 12h of co-culture at two different E:T ratios, 5:1 and 1:1, in the presence 50 pM TcEs and fluorescently labelled Annexin V to monitor cancer cell killing dynamics. Robust T cell-mediated cell death was detected after ∼4 h across all conditions, continuously progressing throughout the experiment, with significant differences in Her2^Low^ cells following the order A>B≥C≥D (**Fig. 3C and D, Supplementary Fig. 3F, G and H and Movies S5 & S6**).

**Figure 3.**
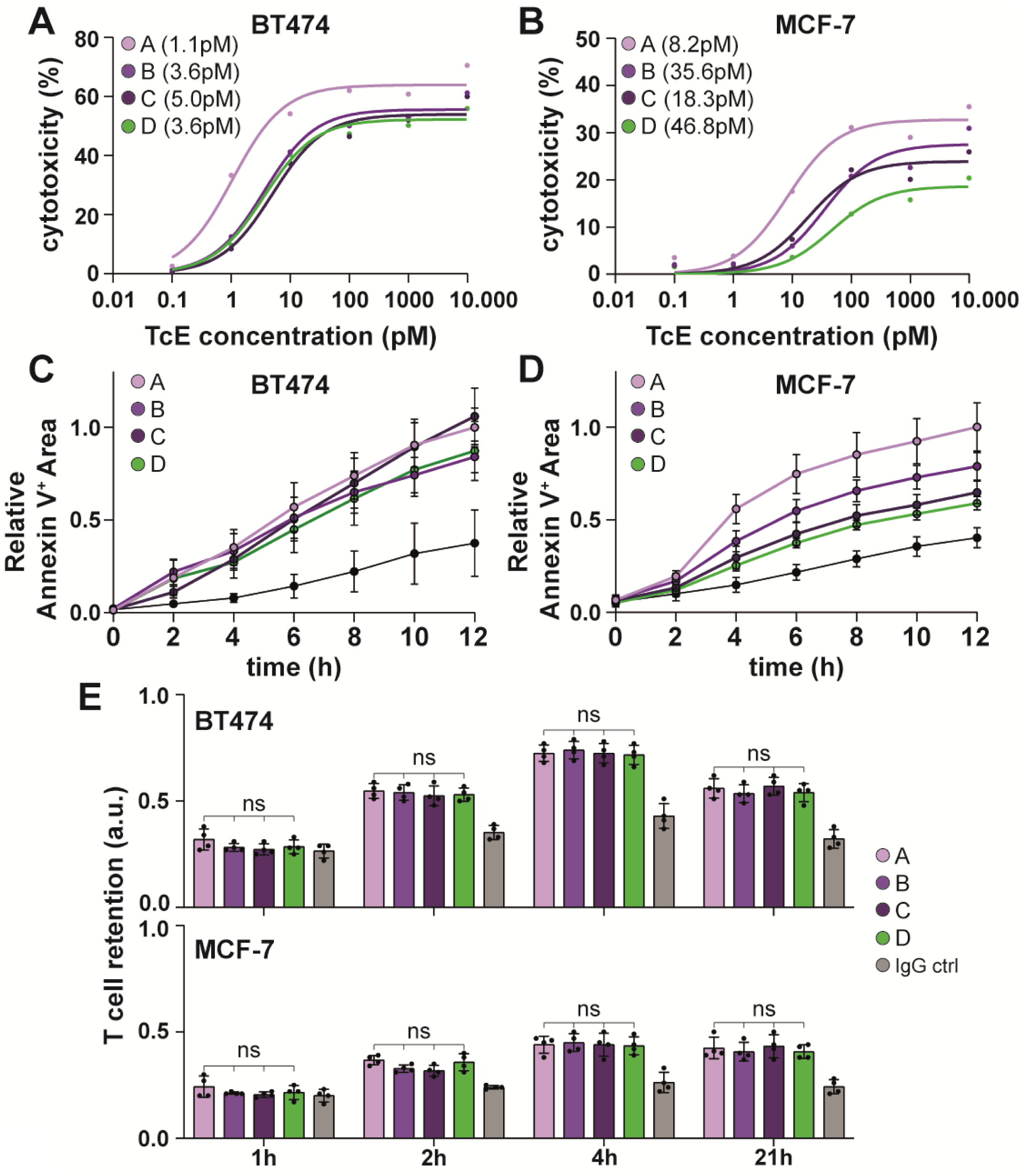
TcE-mediated cytotoxicity and adhesion. (**A**) BT474 or MCF-7 (**B**) cancer cell cytotoxicity upon increasing TcE concentrations after 24h of co-incubation in an E:T ratio of 5:1. Single data points represent the median of 6 biological replicates. Lines represent the non-linear fit of the data. Numbers in brackets = EC50. (**C**) Relative Annexin 5^+^ Area over 12 h in co-cultures of T cells with BT474 or MCF-7 (**D**) cancer cells at an E:T ratio of 5:1 in the presence of 50pM TcEs. 3 biological replicates. Solid circles = Median ± SEM. (**E**) Time-series of the relative retention of T cells on BT474 (top) or MCF-7 (bottom) tumour cells in the presence of 500 pM TcEs or IgG control. Mean ± SD, 2 biological replicates, 2way ANOVA, ns=not significant.

Analysis of early cell-surface T cell activation markers CD69 and CD25 in the context of Her2^High^ cells showed that a 10-fold higher concentration of Format D was required to induce robust upregulation (**Supplementary Fig. 3I**). CD107a cell surface-exposure, a marker for T cell degranulation, which is a key step in T cell mediated cytotoxicity (43), was significantly lower for Format D compared to Formats A-C (**Supplementary Fig. 3J**), especially for Her2^High^ cells. While we did not detect any significant differences in IL-2 and TNFα secretion in response to Her2^High^ cells, Her2^Low^ cells triggered twice as much relative cytokine release with Format A compared to Formats B and C, and no detectable cytokine release with Format D (**Supplementary Fig. 3K**). Collectively, these findings demonstrate that, while all TcEs can induce cytotoxicity and cytokine release, differences between formats exist, especially at low TAA expression levels and can be summarized as A>B=C>D. Next, we aimed to investigate the mechanisms underlying these TcE-format dependent effects.

Initially, we assessed if the different potencies of Formats A-D could be explained by differences in TcE-cell binding. CD8^+^ T cells, Her2^High^ or Her2^Low^ cells were stained with 500 pM TcE and each cell type was washed, incubated with fluorescently labelled secondary antibodies and analyzed by confocal microscopy. No major differences in staining intensity for Formats A-D were observed, indicating that there are no practical differences in TcE-cell binding between the formats (**Supplementary Fig. 4A**).

We then assessed whether relative TcE-potency might be attributed to differences in their ability to mediate cell-cell adhesion. T cells labeled with a fluorescent dye were added to cancer cell monolayers in the presence of 5 pM TcE. Loosely adherent cells were washed away at different time-points, followed by fixation of the cells and recording of the remaining fluorescence intensity. As the tumor cells express ICAM1, CD58 and CD80 (**Supplementary Fig. 3C)**, all of which could contribute to TcE independent adhesion (44), we used human serum IgGs as a negative control for TcE-specific effects. T cell-tumor cell adhesion peaked at approximately 2-4 h of co-incubation but no prominent TcE-format-specific differences in cell-cell coupling could be identified for either cancer cell line (**Fig. 3E**), suggesting that differences in TcE potency are not due to variations in cell-cell adhesion. Thus, we hypothesized that differences in TcE-potency might be attributed to their varying ability to mediate a functional IS.

To test this, we introduced T cells to the SLB system, which we adapted to function as a minimal system for IS formation, by adding ICAM1 at 200 molecules/µm^2^ and mimicking the Her2^High^ and Her2^Low^ expression levels on tumor cells (**Fig. 4A**). TcEs were added at 50 pM in solution, the minimum concentration to induce robust clustering of Her2 in this minimal system while exhibiting pronounced differences in cancer cell cytotoxicity between the different TcEs. Simultaneous addition of T cells and TcEs led to cell adhesion and formation of ISs, whose formation was captured by cell fixation and staining for F-actin. Qualitative inspection of the data revealed striking similarities of TcE-formed adhesion sites to synapses and kinapses observed in natural cell-cell and SLB systems (9), consistent with earlier observations (45). Bulls-eye-like concentric actin structures, reminiscent of dSMACs, were frequently observed (**Figure 4B**), reaching up to ∼50% of total cell contacts for Formats A-C, but only about 30% for Format D, both under Her2^High^ and Her2^Low^ conditions (**Figure 4C**). Her2 accumulated in the center of these structures, similar to the localization of the TCR-pMHC pair in the natural monofocal IS. In this context, the frequency of mature, monofocal IS formation has been associated with the potency of immunotherapies and immune evasion in the tumor microenvironment (46, 47). However, in contrast to natural- or anti-TCR antibody-mediated monofocal synapses, where ICAM1-LFA-1 interactions are observed within a ring-like arrangement in the pSMAC region and excluded from the cSMAC (**Supplementary Fig. 4B**), we found predominantly diffusive and central accumulation of ICAM1 in TcE-mediated adhesions (**Fig. 4B**). Immunostaining for ICAM1 in CD8^+^ T cell and cancer cell co-cultures aligned with this observation, with ICAM1 being enriched but evenly distributed within the cell-cell contact site (**Supplementary Fig. 4C**). Additionally, we assessed the impact of different TcE formats on early TCR signaling, by quantifying the phosphorylation of Zeta-chain-associated protein kinase 70 (pZap-70) at the IS. We found that Formats A-C triggered comparable levels of TCR signaling, while Format D exhibited substantially decreased activity under both Her2^High^ and Her2^Low^ conditions (**Fig. 4D & E**).

**Figure 4.**
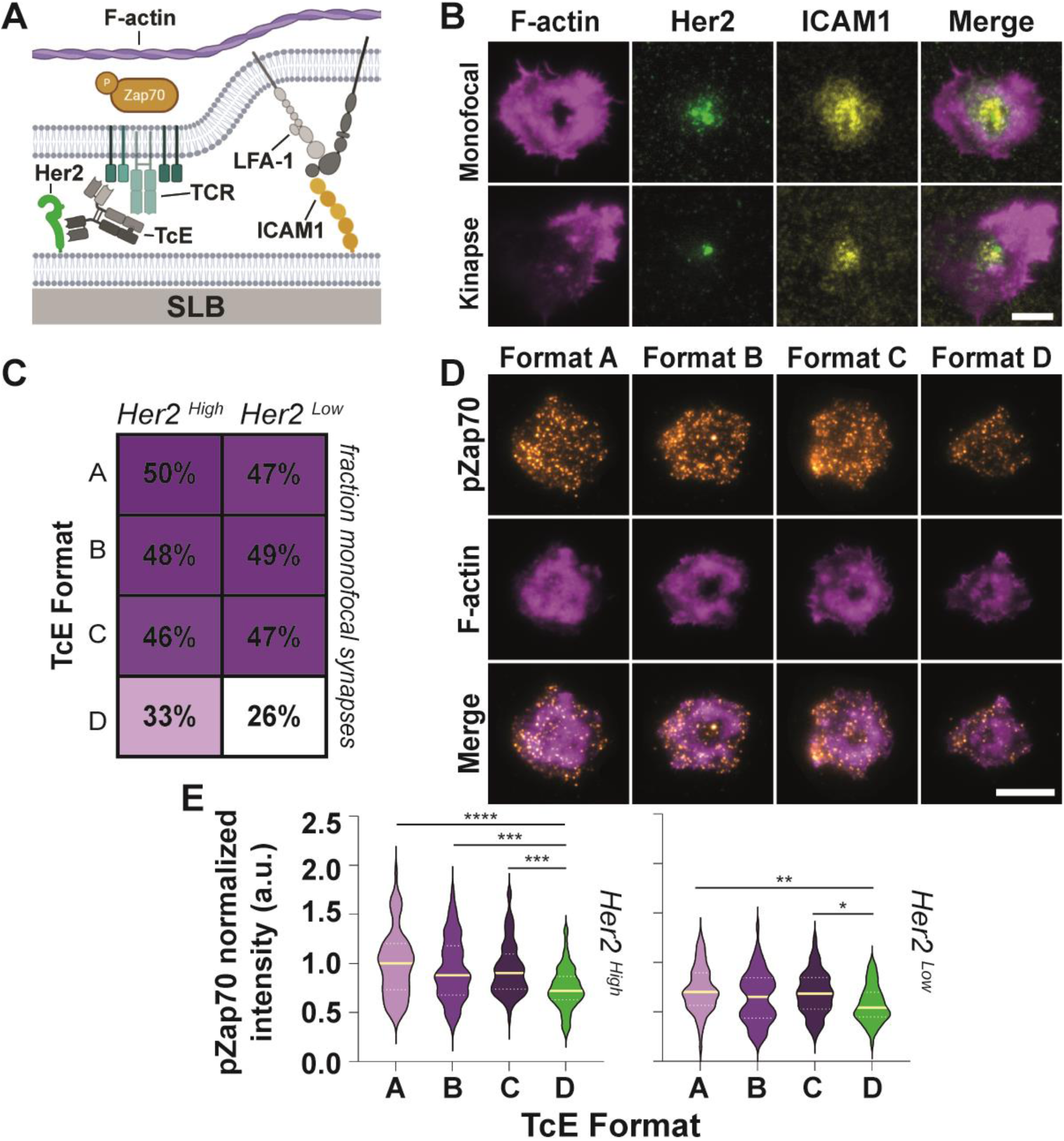
TcE-mediated immune synapse formation dynamics and signaling. (**A**) Schematic overview of T cell-SLB IS experimental system. (**B**) Representative TIRF images of monofocal and kinapse-like TcE-mediated IS formation at 50 pM TcE on SLBs at 400 (Her2, green) and 200 (ICAM1, yellow) molecules/µm^2^ after 20min. of interaction, stained for F-actin (magenta), scale bar = 5µm. (**C**) Analysis of TcE-mediated immune synapse phenotype frequencies after 20min. of interaction. Numbers in boxes represent the percentage of ISs with bulls-eye actin structures. Mean of 3 biological replicates. (**D**) Representative TIRF images of 50 pM TcE-mediated IS formation on SLBs at 400 (Her2) and 200 (ICAM1) molecules/µm^2^ after 20min. of interaction, stained for F-actin (magenta) and pZap70 (orange), scale bar = 5µm. (**E**) Normalized pZap70 levels at the 50 pM TcE-mediated IS at 400 (left) or 30 (right) molecules/µm^2^ of Her2 and 200 molecules/µm^2^ ICAM1. Yellow solid line = median, White dashed line = quartiles. 3 biological replicates, Kruskal-Wallis test, * p<0.05, ** p<0.005, *** p≤0.0007, **** p<0.0001.

Next, we tested the ability of TcEs to recruit co-stimulation at the IS. CD2-CD58 co-stimulation requires close contacts, as noted above, and provides a potent co-stimulatory signal that synergizes with the TCR in the phosphorylation of Phospholipase Cγ1 (pPLCγ1) (11, 48). Notably, the ability to cooperate with the CD2-CD58 interaction has recently been highlighted as an important parameter for chimeric antigen receptors (49) and BiTE-mediated cytotoxicity (50). Addition of T cells to Her2 and CD58 SLBs that had been pre-loaded with TcEs (**Fig. 5A**), followed by colocalization analysis, showed that while all formats had an average positive correlation between Her2 and CD58, this correlation was significantly weaker for Formats C and D, following the trend A=B>C>D (**Fig. 5B & C**).

**Figure 5.**
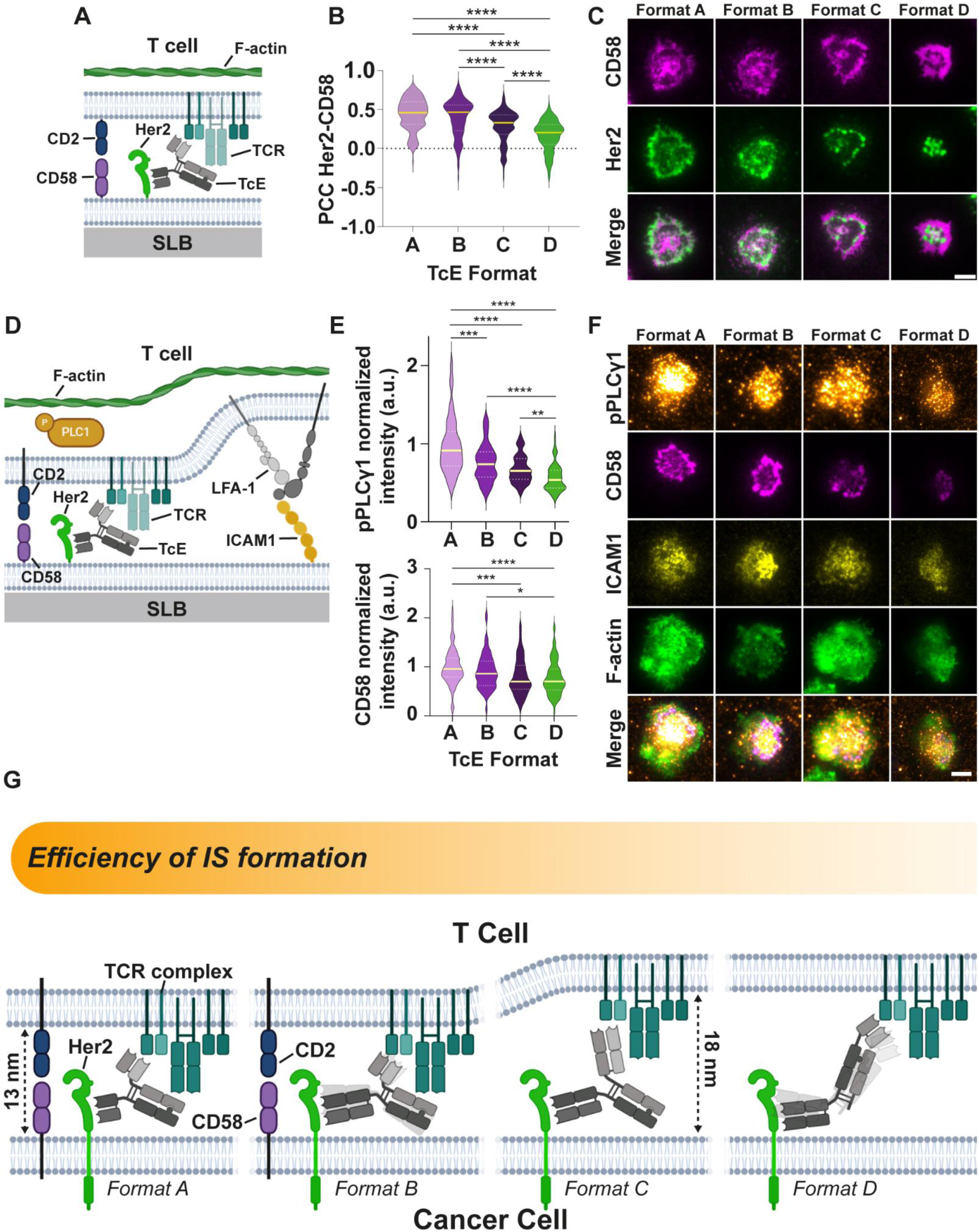
TcE-mediated CD2-CD58 cooperation. (**A**) Schematic overview of T cell-SLB experimental system in the presence of CD58. (**B**) Pearson correlation coefficient of Her2 and CD58 at TcE-mediated T cell-SLB contacts at 400 molecules/µm^2^ of Her2 and 200 molecules/µm^2^ CD58, preloaded with TcEs after 20min. of incubation. Yellow solid line = median, White dashed line = quartiles. 3 biological replicates, Kruskal-Wallis test, **** p<0.0001. (**C**) Representative TIRF images of TcE-mediated T cell-SLB contacts on SLBs from (B) with Her2 (green) and CD58 (magenta), scale bar = 5µm. (**D**) Schematic overview of T cell-SLB IS experimental system in the presence of CD58 and ICAM1. (**E**) Normalized pPLCγ1 (top) and CD58 (bottom) levels at the TcE-mediated IS at 400 (Her2) and 200 (ICAM1) molecules/µm^2^ in the presence of 200 molecules/µm^2^ CD58 and 5pM TcEs after 20min. of incubation. Yellow solid line = median, White dashed line = quartiles. 3 biological replicates, Kruskal-Wallis test, * p≤0.0363, ** p≤0.007, *** p≤0.0007, **** p<0.0001. (**F**) Representative TIRF images of TcE-mediated IS formation on SLBs from (E) with Her2 (green) ICAM1 (yellow), CD58 (magenta), pPLCγ1 (orange) and F-actin (green), scale bar = 5µm. (**G**) Working model of TcE potency. Transparent shadows of Formats B and D represent increased flexibility of the TcE-antigen complex.

Importantly, the addition of CD58 to our Her2/ICAM1 containing minimal system for IS formation revealed pronounced differences in pPLCγ1, which is important for cytoplasmic Ca^2+^ elevation and cytotoxicity. Format A elicited significantly more pPLCγ1 than Formats B and C, which were statistically indistinguishable, followed by Format D with the weakest signaling activity (**Fig. 5D-F**). This was also correlated with the levels of CD58 recruitment to the IS with the order A=B>C=D (**Fig. 5E**). These effects were similar under Her2^Low^ conditions although they were only significant for Format A (**Supplementary Figure 4D**).

Taken together, our results suggest that Format A and B, form CD3ε-TcE-Her2 assemblies that are highly compatible with the ≤13 nm spacing generated by CD2-CD58 interactions, thereby synergistically enhancing pPLCγ1 signaling (**Fig. 5G**). In contrast, bridges formed by Formats C and D with spans of ≥18nm, are less compatible with CD2-CD58 interactions, leading to lower signal amplification and TcE potency. Notably, the greater potency of Format A compared to B, and Format C compared to D, is inversely correlated with their flexibility. This suggests that increased flexibility imposes a penalty on the induction of signaling. Thus, pPLCγ1 signaling emerges as the best and most sensitive predictor for cytotoxicity in our reductionist experimental system as it integrates both close-contact formation and flexibility.

## Discussion

We developed a comprehensive pipeline integrating structural analysis by SAXS, biologically calibrated membrane interface measurements, and IS analysis to assess the potency of IgG-based TcEs. We applied this pipeline to a series of TcEs, varying the spacing between the C-terminus of CD3ε and a membrane proximal epitope of Her2. Our analysis identified key determinants influencing TcE-mediated membrane interface quality, signaling and cytotoxicity at the IS. The first determinant is cell-cell adhesion, a minimal requirement for cytotoxicity, which all TcEs in our series mediated equally well. The second determinant is the formation of close-contacts, which was accomplished by Formats A and B, but not Formats C and D. Close-contact formation with exclusion of large extracellular domains has been previously recognized as important for TcE potency (26, 51, 52). Analysis of SAXS data uniquely revealed a third determinant for TcE potency: flexibility of the CD3ε-TcE-Her2 molecular assembly. This accounted for superiority of Format A over Format B, and superiority of Format C over Format D, as well as equivalence between Formats B and C, as assessed by at least two functional measures for each comparison.

All formats were equally effective at mediating T cell adhesion to cancer cells. Recent studies have suggested that tandem scFv formats can perform poorly on membrane proximal epitopes (27), but Format A, where CD3ε and Her2 targeting scFvs are positioned on different halves of IgG1 Fc, did not face this issue. The employed assay also integrated contributions from adhesion mechanisms, such as LFA-1-ICAM1, which are regulated by TCR signaling that in turn is triggered by TcEs. Format D’s ability to mediate significant adhesion, despite its weaker signaling activity, suggests that it can induce sufficient LFA-1 conformational changes for cell-cell adhesion (53), even based on far-contact, meaning limited CD2-CD58 recruitment, and high flexibility. This sensitive adhesion process creates an interface that allows even the least potent format to achieve significant target killing, even at low TAA expression, establishing the first determinant of TcE quality.

The second determinant, formation of close-contacts (≤13 nm), requires control of intermembrane spacing by the TcE (35, 54). However, we found that simple, linear models of paratope-spacing fail to capture the actual complexity of CD3ε-TcE-Her2 molecular assemblies. 3D measurements from SAXS analysis, combined with CD2-CD58 interactions employed as a molecular ruler (12.8 nm), and Qdot exclusion experiments, revealed that our TcE series can be subdivided into two distinct groups: Formats A and B, mediating a median anchor-to-anchor spacing of ≤13 nm and, and Formats C and D, which mediate a median spacing of ≥18 nm. Formats A and B outperform C and D in the recruitment of CD2-CD58 interactions to sites of Her2 engagement. Notably, however, Formats C and D are able to co-exist with CD2-CD58 mediated close-contacts albeit with significantly lower efficiency, suggesting that even large formats can be compressed into the 12.8 nm gap formed by CD2-CD58. Notably, the role of CD2-CD58 interactions differs in the context of high-affinity TcEs compared to their likely contribution at physiological synapses with rare peptide-MHC complexes. CD2-CD58 is thought to increase sensitivity by facilitating the formation of small, initial close-contacts, allowing the physiological TCR scanning of pMHC (21), as has been observed (55). However, in the case of high affinity TcEs, which bind with orders of magnitude higher affinity than CD2 to CD58 and with more abundant ligands than pMHC, CD3ε-TcE-Her2 assemblies promoted or prevented CD2-CD58 interactions, influencing pPLCγ1 signaling. Our overall results align with an important contribution of the KS model of TCR triggering (21) in that Formats A and B, which mediated close-contacts, were the most potent in the recruitment of CD2-CD58 co-stimulation.

The third determinant of TcE potency that we identified is flexibility of the CD3ε-TcE-Her2 assembly. Flexibility was assessed by SAXS through both information theory-based analysis of multistate modeling results and independently through Kratky plots. We speculate that flexibility influenced the potency of TcEs to induce intrinsic TCR-triggering events, such as greater mechanical coupling leading to TCR conformational changes. Greater flexibility of close contact forming Format B may account for its lower pPLCγ1 signaling activity compared to Format A. Furthermore, lower flexibility may account for the ability of far-contact forming Format C to trigger similar pPLCγ1 signaling as close-contact forming Format B, with nearly identical cytotoxic and cytokine release potency. A similar flexibility penalty was also noted in the analysis of agonistic anti-CD40 antibodies bearing mutations in the hinge (56). Additionally, less flexible molecules tend to exhibit an increased half-life at membrane-to-membrane interfaces compared to in-solution measurements, leading to enhanced signaling (57, 58). Flexibility has been seen as a desirable feature when it promotes binding by the TcE, which we would relate to the adhesion requirement (27). Flexibility emerged as an interesting parameter in our analysis, but this was an unexpected opportunity that emerged from the SAXS analysis and was not planned at the design stage. Future research should extend the analysis of TcE flexibility, for example, through systematic manipulation of the hinge region (56), to further isolate flexibility as a parameter in TcE design. Notably, other features that varied across the TcE formats were lower thermal stability and higher hydrophobicity associated with the scFvs, following the trend A>D=B>C (**Supplementary Figure 1C**). The high thermal stability and low hydrophobicity of Format C may account for selection of similar IgG1-like configurations for the majority of clinically approved TcEs.

Our findings have significant implications for understanding the function of existing clinically approved TcEs and those in clinical trials. The structure of BiTE formats like Blinatumomab (59), the first approved TcE, or recently approved Tarlatamab (60), suggest that the CD3ε-BiTE-TAA complex may form close contacts with more membrane distal epitopes, such as on CD19, with minimal flexion, leading to high potency. However, many other approved TcEs target membrane-proximal epitopes such as CD20 (61). Our pipeline could be applied to the analysis of Format C-like TcEs like Epcoritamab (62), Mosunetuzamab (63) and Odronextamab (64), as well as BCMA-targeting Teclistamab (65), potentially even using detergent micelles or nanodiscs. Plamotamab (66) is a Format B-like TcE, which we would predict to perform equally to Format C-like TcEs assuming comparable CD2-CD58 recruitment and higher flexibility. Glofitamab, on the other hand, is Format C-like, but has an additional anti-CD20 paratope linked to the Fab arm bearing the anti-CD3 (67), potentially reducing the flexibility of the TcE within the synapse. It would be important to determine which of these effective drugs fully utilizes CD2-CD58 or other similarly sized co-stimulatory molecules. There are no Format D-like TcEs approved, but its low cytokine production may be beneficial in settings where cytokine release syndrome limits the efficacy of therapy.

Finally, we acknowledge that our SLB system does not fully recapitulate the active participation (e.g. F-actin based transport and ligand immobilization), potential biophysical cues (e.g. membrane topology, elasticity, etc.) and full repertoire of co-stimulatory signals and glycocalyx components of the target cell. However, it has been shown that IS function and architecture in SLBs are well correlated with natural ISs formed at cell-cell contacts and therefore represents a realistic model system that allows accurate measurements (12). Importantly, our SLB system can be easily adapted to present different TAAs, additional co-stimulatory ligands and other components of the glycocalyx (55), which will make it an invaluable tool for the early characterization of new TcE designs against various surrogate cell-surface compositions.

In summary, we have established a pipeline including SAXS and SLB-based experiments for TcE analysis that enables effective prediction and interpretation of TcE performance. Our data identified three key determinants of TcE quality for a membrane proximal TAA: 1.) cell-cell adhesion, 2.) close-contact formation and 3.) low flexibility of the CD3ε-TcE-TAA assembly. We suggest that consideration of these determinants maybe useful in design and evaluation of future TcE formats, allowing precise tuning of their performance.

## Material and Methods

### TcE design

Genes encoding variants of anti-Her2 Fabs and anti-huCD3ε (I2C, Amgen) were obtained by gene synthesis of light chain and heavy chain variable domains (Genewiz). Human IgG1 Fc with effector function knock-out (LALA) mutations was used as Fc-region for all TcEs, which additionally contained Genentech’s knob-in-hole mutations in the CH3 domain to promote heterodimerization (68).

### TcE expression and purification

TcEs were isolated from CHO cells that were transfected with respective expression plasmids. Supernatants were harvested 7 days after transfection, followed by sterile filtration through a 0.2 µm filter. TcEs were isolated in a two-step process using protein A chromatography, followed by size exclusion chromatography. Eluted fractions were verified by SDS-PAGE.

### Small-angle X-ray scattering (SAXS)

TcE-antigen complexes were prepared by mixing TcEs with a 1.3 molar excess of Her2 (ACROBiosystems, HE2-H5225) and CD3εγ, containing ∼30 amino acid dimerization domains on each monomer (ACROBiosystems, CDG-H52W5). TcEs and TcE-antigen complexes were frozen and shipped on dry ice to the Diamond Light Source synchrotron, Beamline B21 (69) for SAXS analysis. The software ScÅtterIV (developed by Robert Rambo, Diamond Light Source, Oxford, UK) was used for initial manual inspection of the SAXS curves and removal of outliers, resulting in a single, optimized raw SAXS data curve for every TcE or TcE-antigen complex. Multi-state modelling was performed with MultiFoXS (70), based on existing start models from the protein data bank (PDB) or AlphaFold2 (71). Single domains and linkers were assembled in UCSF Chimera (72) and Coot (73). Differences between the calculated and experimental scattering intensities were measured with MultiFoXS to determine the quality of the models, resulting in 60-156 states per TcE or TcE-antigen complex. Paratope (distances between the tips of the heavy chain CDR3 loops) and anchor-to-anchor spacings (distances between the C-terminal domains of CD3εγ and Her2) and measures of flexibility were calculated using custom Text Command Language (TCL) codes in VMD (74), taking into consideration the weighted effective contribution of each PDB file. Dimensionless Kratky plots were directly calculated from the optimized SAXS curves.

### Supported lipid bilayers (SLBs)

SLB preparation has been described in detail elsewhere (75). In brief, SLBs were generated from small unilamellar vesicles prepared by mechanical extrusion containing 12.5 mol% DOGS-NTA(Ni^2+^) and 87.5 mol% DOPC (all lipid reagents obtained from Avanti Polar Lipids, USA) used at a final lipid concentration of 0.4 mM. Plasma-cleaned glass cover slips (SCHOTT UK Ldt, UK) were fixed to 6-channel flow chambers (sticky-Slide VI 0.4, Ibidi, Thistle Scientific LTD, UK). Each channel was filled liposome solution and incubated for 30 min. at RT. Each channel was washed twice with flow cell buffer (20 mM HEPES pH 7.2, 1.37 mM NaCl, 5 mM KCl, 0.5 mM Na_2_HPO_4_, 6 mM D-Glucose, 1mM CaCl_2_, 2 mM MgCl_2,_ 1% BSA). SLBs were blocked with 3% BSA in flow cell buffer for 20 min., followed by two washes and subsequent incubation with His-tagged and fluorescently labelled recombinant Her2 for 60 min., adjusted to a concentration to obtain 30 or 400 molecules/µm^2^. For some experiments SLBs were additionally coated with 200 molecules/µm^2^ recombinant human AlexaFluor405-ICAM1-12His or 200 molecules/µm^2^ AlexaFluor647-CD58-12His. For experiments with Quantum Dots, 0.001% CapBio lipids were added during SLB formation. Streptavidin coupled QDot525 (Thermofisher) was added at 50nM. Specific protein concentrations needed to obtain the desired molecular densities on the SLBs were deduced from flow cytometry-based calibration experiments on bead-supported lipid bilayers compared to fluorescently-labelled reference beads of know molecular densities (Bangs Laboratories, IN). For synapse formation experiments, CD8^+^ T cells were injected into the flow channels to reach a final concentration of 100,000 cells/channel. Prior to injection into the channels, cells were mixed with TcEs to reach an end concentration of 5-50pM. For some experiments, SLBs were pre-loaded with TcEs at a concentration of 10nM, followed by three washes with wash buffer before cells were introduced. Cells were allowed to interact with the SLBs for the respective times in a cell-culture incubator at 37°C, 5% CO_2_, followed by with 4% PFA for 20 min. Cells were permeabilized with 0.1% Triton-X100 for 2 min, blocked with 1% BSA in PBS for 20 min. and, where required, subsequently stained with fluorescently labelled Phalloidin and anti-human IgG secondary antibodies for 1 hours. To stain for pZap70 and pPLCγ1, cells were blocked for 1h with 5% BSA in PBS and then either stained with rabbit anti-pPLCγ1 (Cell Signaling, #2821, 1/100) or rabbit anti-pZap70 (Cell Signaling, #2704S, 1/100) overnight at 4°C. Samples were then washed three times with PBS and stained with goat anti-rabbit AlexaFluor Plus 405 secondary antibodies (Thermofisher, A48254, 1/300) for 1h at room temperature. Imaging was performed on a Olympus IX83 inverted microscope (Keymed, Southend-on-Sea, UK) equipped with 405-nm, 488-nm, 561-nm and 640-nm laser lines, a Andor sCMOS Prime 95B camera and a 100x 1.45 NA oil-immersion objective.

### Giant unilamellar vesicle (GUV) preparation

GUVs were prepared by mixing small unilamellar vesicles (1mol% DGS-NTA(Ni^2+^), 1mol% PE-Lissamine Rhodamine, 20 mol% EggPG, 78 mol% EggPC), containing 10mM MgCl_2_, with a perfluorinated oil phase, containing 1% (w/w) stabilizing non-ionic surfactant and 10 mM of an anionic PFPE-COOH surfactant in a 1:2 ratio (76). Manual shaking was used to generate an emulsion that formed droplet-supported GUVs, which were released by addition of 1H,1H,2H,2H-Perfluor-1-octanol. GUVs were loaded with CD3εδ heterodimers, containing ∼30 amino acid dimerization domains on each monomer (Acro Biosystems, CDD-H52W1), and fluorescently labelled, CD2-10His. GUVs were added to SLBs for 1h, followed by image acquisition.

### Statistics and data analysis

All statistical analysis, linear regressions and plotting were performed with GraphPad Prism 9.

## Acknowledgments and funding sources

Kennedy Trust for Rheumatology Research and Boehringer Ingelheim Pharmaceuticals. A.L. was funded by an Erwin Schrödinger postdoctoral fellowship of the Austrian Science Fund (FWF, project number: J4542-B) and was an EMBO non-stipendiary postdoctoral fellow (ALTF 1109-2020). O.S. acknowledges support from the Joachim Herz foundation (Add-on fellowship). A.L. and M.D. are supported by the Biotechnology & Biological Sciences Research Council (BBSRC, BB/X015408/1). T.M. was supported by the CAMS-Oxford Institute Senior Postdoctoral Research Fellowship, funded by the CAMS Innovation Fund for Medical Science (CIFMS), China (Grant: 2018-I2M-2-002). O.D. is funded by a Wellcome Trust Senior Fellowship in Basic Biomedical Sciences (207537/Z/Z17/Z). We would like to thank the antibody expression and purification (APEX) team at Boehringer Ingelheim for providing high quality material to support this study. The authors thank Claire Pizzey and her dedicated team at the synchrotron beamline B21 at Diamond Light Source (Oxford, UK) for their organization and execution of the SAXS measurements, as well as their valuable advisory support.

