## Supporting Information for "Solution structure and synaptic analyses reveal determinants of bispecific T cell engager potency"

* contributed equally
\# contributed equally

**This PDF file includes:**

SI Material and Methods
Figures S1 to S4
Legends for Movies S1 to S6
SI References

**Other supporting materials for this manuscript include the following:**

Movies S1 to S6

### SI Material and Methods

***Analytical Size-exclusion chromatography (aSEC):*** aSEC was carried out using the Agilent HPLC system. aSEC, in combination with multi-angular light scattering (MALS) was used to determine the molecular weight and quality of TcEs.

***Differential Scanning Calorimetry (DSC):*** DSC was carried out on a MicroCal VP-Capillary calorimeter (Malvern, UK) equipped with an autosampler, following previously published methods (1).

***Analytical Hydrophobic Interaction Chromatography (aHIC):*** Samples were mixed 1:1 with 2 M $(NH_4)_2SO_4$ and analyzed using a Waters Acquity UPLC H-class system with a Sepax Proteomic HIC butyl-NP 1.7 column. Chromatograms were generated from absorbance at 280 nm as a function of decreasing $(NH_4)_2SO_4$ concentration. Data were analyzed using Empower 3 software (Waters).

***Surface plasmon resonance (SPR):*** Experiments were performed on a Biacore T200 instrument (Cytiva) at 37°C. Human CD3ε and Her2 (SinoBiological, 10004-H08H and 10977-H08S) were amine-coupled to a CM5 chip (Cytiva) at ca. 280 response units using an amine coupling kit (Cytiva). A 3-fold dilution series for each TcE was injected and sensograms were recorded with the following parameters: association time: 2 min; dissociation time: 25 min; flow rate, 100 µl/min. After each cycle, the surface was regenerated by injecting 3 M $MgCl_2$ to fully unbind bound TcEs. Data were analyzed with GraphPad Prism using the 'Association then dissociation' function with a global fit for $k_{ON}$, $k_{OFF}$, and $B_{max}$.

***Small-angle X-ray scattering:***

*Sample preparation and data acquisition:* Amicon Ultra centrifugal filter units with a molecular weight cutoff of 30 kDa (Merck Millipore) were used for buffer exchange to 50 mM Tris 150 mM NaCl pH 7.5, 2% glycerol and subsequent concentration of TcE samples. TcE samples without antigens were concentrated to 1, 2, 4 and 10 mg/ml and 50 µL per concentration were used for SAXS measurements. TcE-antigen complexes were prepared by mixing TcE molecules with 1.3 molar excess of Her2 ectodomain and CD3εγ (ACROBiosystems, HE2-H5225 and CDG-H52W5), respectively. After 1h of incubation on ice, the buffer was subsequently exchanged and samples were concentrated to 10 mg/ml. Final sample volume for measurement was 50 µL. Samples were frozen and shipped on dry ice. SAXS measurements were performed at the Diamond Light Source synchrotron, beamline B21. TcE samples were measured in batch mode, while TcE-antigen samples were measured by inline SEC-SAXS. Batch mode samples were handled by an Arinax BIOSAXS liquid sample-handling robot. SEC-SAXS samples were injected into a Superdex 200 column (Cytiva) and run on an inline Agilent 1200 HPLC system. Data was collected using an EigerX 4M detector (Dectris, CH) at a sample-detector distance of 3689 mm and a wavelength (λ) of 0.95 Å. The B21 beamline was operated at 13.0 keV. The covered range of momentum transfer (scattering vector q) was $0.0045 < q < 0.34$ Å$^{-1}$ ($q = 4\pi * \sin(\theta)/\lambda$, where θ is the scattering angle).

*Initial data processing:* Raw SAXS curves were initially processed using the software ScÅtterIV (developed by Robert Rambo, Diamond Light Source, Oxford, UK). The baseline corrected curves of the different TcE concentrations were manually inspected, and small scattering vector values of high concentration samples were discarded if signs of aggregation were visible. Similarly, large scattering vector values of low concentration samples were discarded if necessary, depending on background noise, yielding improved SAXS curves through combining measurements at different concentrations (2). Peak regions of the TcE-antigen complexes were selected in ScÅtter, based on signal intensity and similarity of software calculated $R_g$ values. Measuring points with similar $R_g$ values in the range of the main intensity peak were selected and merged to a single SAXS curve. In total, these procedures resulted in a single, optimized raw SAXS data curve for every TcE or TcE-antigen complex.

*Multi-state modelling:* For further processing, optimal scattering vector ranges were selected from SAXS curves: small scattering vector values, lacking linearity in Guinier Peak analysis, and large scattering vector values with high intensity fluctuations, were discarded. Optimal ranges of the SAXS curves were employed for multi-state modelling using MultiFoXS. Molecular start models were generated based on existing protein data bank (PDB) models (PDB entries: 1N8Z (Her2), 1SY6 (CD3εγ), 1IGT + 1HZH (IgG)) and AlphaFold2. Single domains and linkers were assembled in UCSF Chimera and Coot. Flexible linkers, connecting heavy and light chains of scFvs and scFabs were omitted. Conformational sampling of the start models was done by using the rapidly exploring random tree (RRT) algorithm of the Integrative Modeling Platform (3). A pool of 10,000 conformations (saved as PDB model files) per structure were generated.

Unstructured linker residues of the hinge region, as well as the linker between the C-term of the Fc part and the anti-CD3-Fv of Format D, were defined as flexible, while keeping the connectivity of the disulfide bonds in the hinge region. For the TcE-antigen models, the unstructured N-terminal antibody-binding motif of CD3ε was additionally defined as flexible. All other parts of the models were treated as rigid bodies. For every conformation, the theoretical SAXS profile was pre-computed using the “foxs” command. Finally, multi-state models were generated with the “multi_foxs” command, using the pre-computed conformation profiles and the experimental SAXS data. The output was ensembles of conformations (PDB structures) of different sizes (1-6). Each ensemble fitted the experimental SAXS curve, quality of fits was determined by MultiFoXS, using the differences between the experimental and the calculated scattering intensities, expressed by the chi-squared value of the ensembles. The 10 best-scoring ensembles (chi-squared value range: 0.01 - 1.91 for TcE models; 0.88 – 18.57 for TcE-antigen models) of every ensemble size were used for further analysis. Based on the algorithm-determined maximal ensemble sizes, this resulted in 60-156 states (molecular PDB models) per TcE or TcE complex, representing the conformational space that is covered by the respective molecule or complex in solution.

*Effective weights of unique PDBs, PDB alignments, and weighted statistics*: Each ensemble was generated independently, allowing certain PDB structures (states) to recur across multiple ensembles, indicating their dominance in the conformational space. Within each ensemble, the intra-ensemble weights of the states reflect their relative frequencies. Consequently, the same state may appear in different ensembles with distinct weights, reflecting its specific contribution within the unique composition of each ensemble. Assuming equal likelihood of each ensemble, effective weights for unique states (PDB conformations) were calculated by aggregating their intra-ensemble weights and dividing by the total number of ensembles. This normalization accurately represents the contributions of unique states across ensembles, collectively defining the conformational space. The quantitative analyses as depicted by the violin plots for paratope distances, radius of gyration, flexibility of center of masses and rotational flexibility, were performed over the unique PDB conformations using custom Text Command Language (TCL) codes and MATLAB, weighted by their effective contributions. The PDB structures were aligned to either anti-Her2 in complex with Her2 (for TcE-antigen complexes) or anti-Her2 (for TcEs)

*Distance measurements, flexibility analysis and visual representation:* For paratope spacing analysis, the direct distances between the tips of the heavy chain CDR3 loops for every single state were measured (custom TCL codes). For TcE-antigen complexes the direct distances between the C-terminal domain of CD3ε and the C-terminal domain of Her2, respectively (custom TCL codes). Measurements were weighted according to the effective weighting of states.

The flexibility of center of mass (CoM Flex) of a molecular domain or the entire molecular assembly corresponding to a state (PDB conformation) was defined as the distance between the weighted average center of mass (WACoM) of the molecular domain or the entire molecular assembly and the center of mass (CoM) of the same domain or assembly in the given conformation.

The rotational flexibility of a molecular domain or the entire molecular assembly corresponding to a state (PDB conformation) was quantified as the angle between two axes: (1) the axis connecting the WACoM of HER2 (for TcE-antigen complexes) or anti-HER2 (for TcE) to the WACoM of the relevant domain or whole assembly, and (2) the axis connecting the WACoM of HER2 (for TcE-antigen complexes) or anti-HER2 (for TcE) to the CoM of the same domain or assembly in the given conformation.

To quantify the Shannon entropy of a molecular domain or the entire molecular assembly, we utilized the product of 'CoM Flex' and rotational flexibility, weighted by the state-specific distributions. First, the product values were binned into a manually defined histogram, with bin edges determined by the range of values divided into a specified number of bins. Weighted counts for each bin were calculated by summing the weights of values falling into each bin. These weighted counts were normalized to obtain probabilities, which were used to compute the Shannon entropy as the negative sum of the weighted probabilities multiplied by their natural logarithms.

Dimensionless Kratky plots were directly calculated from the optimized SAXS curves ($I(q)/I(0)*(q*R_g)^2$ vs. $q*R_g$) and plotted with GraphPad Prism. $R_g$ values were determined from the SAXS curves using the program Primus from the ATSAS software suite. $R_g$ and $D_{max}$ values of the single states (PDB models) of the multi-state model ensembles were determined by the program FFMAKER from the ATSAS software suite.

Visual representations of the conformational space as presented in the main text (Figure 1B) were created in UCSF Chimera. For these representations, all PDB models of the 10 best-scoring ensembles of every ensemble size were aligned to the Fc part. The envelope shape of all conformations of a TcE molecule or complex was created by using the “molmap” command on all PDBs, rendered at 20 Å resolution. The envelop surface was displayed in mesh representation. For visual guidance, the model of a single state was selected and displayed in surface representation. Visual representations of the conformational space of TcE-antigen complexes as presented in Supplementary Figure 2F and Movies S1-S4 were was created in VMD using customized TCL codes. For these representations, the unique PDB models were aligned to anti-Her2 in complex with Her2 and depicted with weighted lines in accordance with their effective weights in the conformational space. While the representations were rendered using TachyonInternal, the videos were generated using VideoMach.

*Weighted polar histograms of tilt angles relative to the Her2–anti-Her2 axis*: To analyze the orientations of molecular domains or entire molecules within the conformational space, weighted polar histograms were employed, as depicted in Figure SX. These histograms capture the tilt angle distributions for each defined domain (e.g., Fc, anti-CD3) relative to the Her2–anti-Her2 axis, designated as the positive z-axis in a locally defined right-handed coordinate system. The polar angles in the histograms (ignoring the sign) represent the angular displacement from the positive z-axis, ranging from 0 to 180 degrees. The angles were assigned a negative sign if the center of mass of the domain had a negative y-component in the local right-handed coordinate system, with the Her2 WACoM as the origin and the Her2-to-anti-Her2 WACoM axis defining the positive z-direction.

***Cell culture:*** Human primary $CD8^+$ cells and MCF-7 and BT747 cancer cell lines (obtained from the American Tissue Culture Collection) were routinely cultured in RPMI (Gibco) medium supplemented with 10% fetal bovine serum, 1% penicillin/streptomycin, 1% L-glutamine, 1% non-essential amino acids and 50 mM HEPES in cell culture treated flasks at 37°C, 5% $CO_2$ and 100% humidity. Cancer cells were split at ~80% confluency by trypsinization. For $CD8^+$ culture and expansion as well as for all functional assays described, 50 U/mL recombinant human IL-2 (PeproTech, UK) were added to the culture medium. Primary human $CD8^+$ were isolated from healthy donor blood, provided by the National Health Service blood service under ethics agreement number 11/H711/7, with commercially available negative selection kits (RosetteSep Human $CD8^+$ T cell Enrichment Cocktail, STEMCELL technologies, UK) following the manufacturers instruction. After isolation, cells were expanded by addition of anti-CD3/anti-CD28 T-cell activation beads (Dynabeads ThermoFisher Scientific, UK) at 25 µL/ $10^6$ cells for 3 days following the manufacturer's instructions. The magnetic beads were subsequently removed and cells were expanded further at $10^6$ cells/mL for an additional 4 days. T cells were either used within 2 days for experiment or cryopreserved.

***Cell staining and immunofluorescence:*** Staining of cells with CellTracker Green CMFDA and CellTracker Red CMTPX (Thermo Fischer Scientific, USA) was performed at a final concentration of 5 µM for 30 min at 37°C. For quantification of surface exposure of degranulation marker Lamp-1, human primary T-cells were cocultured with cancer cells as described in the "Functional Assays" section. After incubation for 21 hours, cultures were resuspended 10 times by pipetting up and down. 100 µL of the culture medium, containing T cells, were then collected and fixed for 20 min. by dropwise addition of PFA to an end concentration of 4%. Cells were then washed 2x with PBS and blocked with 1% BSA for 30 min. No permeabilization was performed. Cells were then stained with monoclonal mouse anti-human Lamp-1-AlexaFluor488 (BioLegend 328610, clone H4A3) at a final dilution of 1:500 for 1 hour. Excess antibody was removed by washing twice with 1% BSA in PBS. Samples were recorded on a BD Fortessa X20 machine equipped with an HTS sampler. LAMP-1 staining was quantified by pre-gating on CellTracker Red stained T cells.

***Confocal imaging, TIRF imaging and image analysis:*** Confocal microscopy imaging was performed with a LSM880 laser scanning confocal microscope, equipped with a Plan-Apochromat, 63X, numerical aperture 1.4 oil objective. TIRF imaging was performed with an Olympus IX83 inverted microscope (Keymed, Southend-on-Sea, UK) equipped with 405-nm, 488-nm, 561-nm and 640-nm laser lines and a Photomertrics Evolve delta EMCCD camera. Images were acquired with a 150x 1.45 NA oil-immersion objective. If not specified otherwise, a minimum of 30 images was recorded for each sample for each biological replicate.

All image post-processing and analysis was performed with ImageJ (NIH). Adjustments to image brightness and contrast, as well as background corrections, were always performed on the whole image and special care was taken not to obscure or eliminate any information from the original image. For semi-automated quantification of signal intensities, all images were first background-correct with a 50-pixel rolling ball algorithm and converted to 8-bit images. Cell attachment sites were segmented based on the actin staining using the Ilastik (4) machine-learning based toolkit. The resulting cell masks were then used to quantify the fluorescence signal in the channel of interest using ImageJ's 'Analyze Particles' function. For quantification of synapse frequencies with bulls-eye structure, cell masks were used to quantify the total number of cells per sample and the fraction of cells with an actin-depleted region in the center, corresponding to a monofocal synapse.

***TcE-cell binding analysis****:* For analysis and quantification of TcE binding to $CD8^+$ T-cells, MFC-7 and BT474 cultures, cells were treated with a trypsin solution for the minimum time required to release them form the surface followed by trypsin-inhibition with serum containing media, washing with PBS and dilution to a final concentration of $10^6$ cells/mL. Cells were then incubated with TcEs diluted in fully-supplemented culture medium (5 pM final TcE concentration) medium for 1 hour and subsequently washed twice with fully-supplemented culture medium. Goat anti-human IgG-PE (Thermo Fischer, 12-4998-82) was then added to the cells at a final dilution of 1:500 for 30 min., followed by washing twice with fully-supplemented culture medium and immediate imaging by confocal microscopy. Mean fluorescence intensities were calculated from maximum intensity projections of z-stacks.

***Functional assays:*** If not specified otherwise, all functional assays involving primary human $CD8^+$ T-cell/ cancer cell cocultures were performed in 96-well transparent flat bottom well plate formats (Costar 96) with a total culture volume of 150 µL and a T cell to cancer cell ratio of 1:5 with a co-incubation time of 24 hours in the presence of 50 U/mL recombinant human IL-2. Cancer cell lines were seeded on the day prior to the experiment in 100 µL fully-supplemented culture medium without recombinant human IL-2 at a concentration of 10,000 cells/well to allow adhesion. Before addition of TcEs and control IgGs, 50,000 $CD8^+$ T-cells were added in 50 µL to each culture well and subsequently TcEs were added without further delay. For time resolved functional assays, multiple technical replicates were prepared. Culture medium and cells were then collected at the indicated time points. If not stated otherwise, final TcE concentration was adjusted 500 pM.

LDH release assays (CyQuant LDH cytotoxicity assay, Thermo Fischer, USA), human IL-2 ELISA (human IL-2 Quantikine ELISA kit, R&D systems, USA) and TNFα ELISA (human TNF-alpha Quantikine ELISA kit, R&D systems, USA) were performed from cell culture supernatants, following the manufacturers´ instructions.

Adhesion assays were performed by seeding cancer cells at 10,000 cells/well in transparent flat bottom 96-well plates in 100 µL, followed by incubation over night to allow for cell adhesion. Subsequently, human primary T-cells, stained with CellTracker Red, were added at a concentration of 50,000 cells/well in 50 µL to the cancer cells and co-incubated for the indicated time periods. After incubation, cells were washed 3x with PBS to remove unbound and loosely-attached T cells. Cultures were then fixed by addition of 100 µL 4% PFA, which was replaced after 20 min. by PBS. Fluorescent intensities of the CellTracker Red signal in each well were then measured in each well with a FluoStar Omega microplate reader in fluorescence mode with a total number of 10 flashes in orbital averaging mode and excitation/emission filters set to 544/590 maxima (gain 2500).

***Live monitoring of cancer cell cytotoxicity***: BT474 or MCF-7 cancer cells were stained with 20µM CFSE, seeded into the wells of an 18-well slide (iBidi) at 20,000 cells/well and incubated over night at 37°C, 5% $CO_2$. T cells were added at 20,000 or 100,000 cells/well (1:1 or 5:1 E:T ratio respectively) in the presence of 50pM TcEs and 1/200 Red Incucyte Annexin-V dye (Sartorius). Imaging was performed on a LSM980 confocal microscope equipped with an absolute focus module in multi-position mode. Images were collected every 30 min. for 12h. Image analysis was performed by segmenting the CFSE and Annexin-V signals using machine learning based image segmentation software Ilastik (4), followed by calculation of the relative Annexin-V positive area in ImageJ.

***ICAM1, CD58 and CD80 immunostaining****:* For analysis of ICAM1, CD58 and CD80 surface expression on cancer cell lines, cells were seeded in 96-well flat bottom transparent well plate formats with a total culture volume of 100 µL for 24 hours with 50 U/mL recombinant human IL-2. Cells were subsequently washed twice with PBS and incubated with AlexaFluor647-anti-human-ICAM-1 (BioXcell, BE0020-2, 3 µM), anti-human AlexaFluor647-anti-human-CD58 (BD Bioscience, 563567, 1:500 dilution) or anti-human AlexaFluor647-anit-human-CD80 (BioLegend, 305216, 1:500 dilution) for 30 min at room temperature. Cultures were then washed twice with fully-supplemented culture medium and fixed with 4% PFA for 20 min followed by washing twice with 1% BSA (in PBS). Staining was evaluated by laser-scanning confocal microscopy. For analysis of ICAM1 localization in $CD8^+$ T-cell/ cancer cell cocultures, cells were seeded and incubated as described in the “functional assays” section and immunostained as described above.

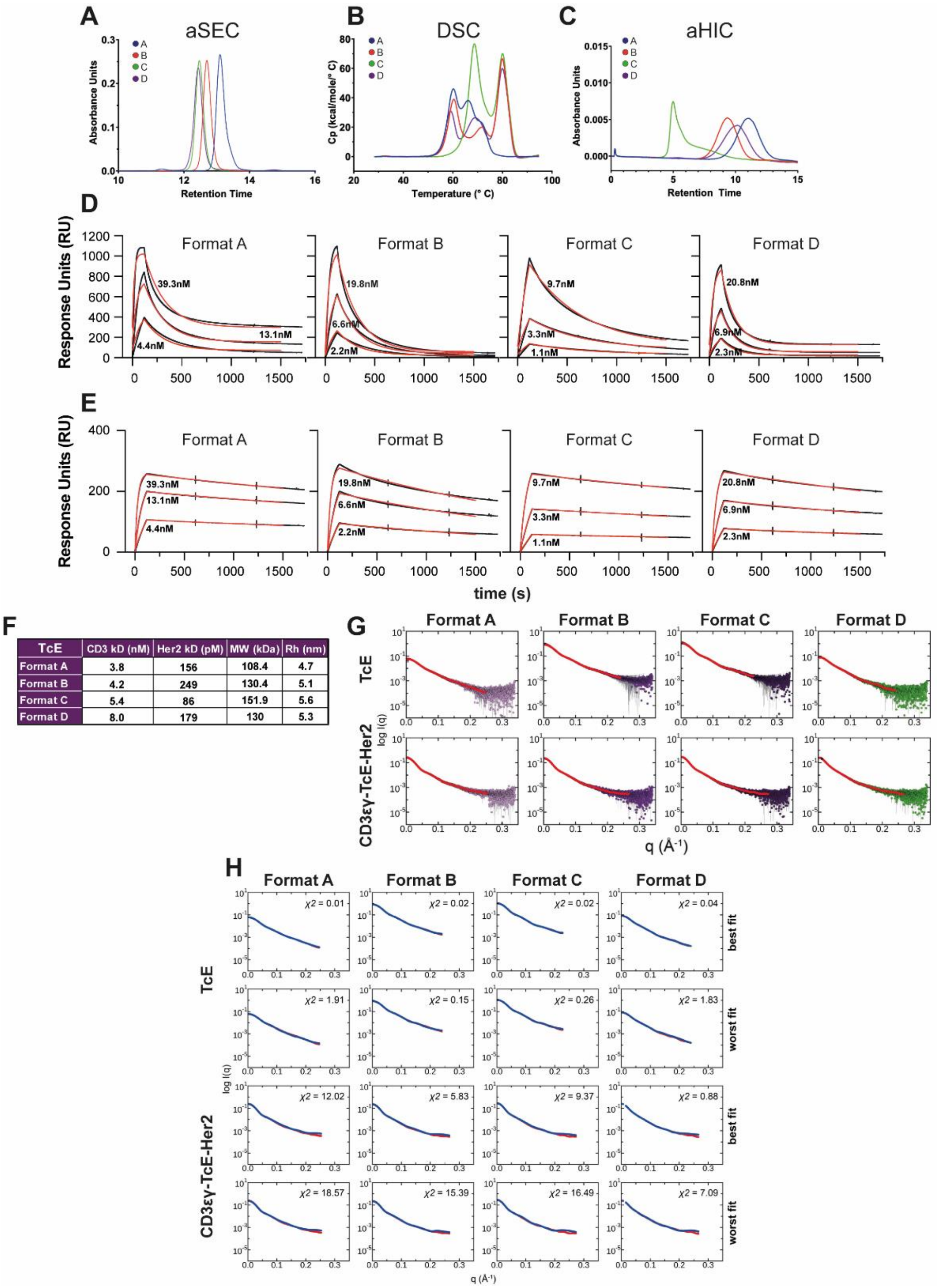


| TcE | CD3 kD (nM) | Her2 kD (pM) | MW (kDa) | Rh (nm) |
|---|---|---|---|---|
| Format A | 3.8 | 156 | 108.4 | 4.7 |
| Format B | 4.2 | 249 | 130.4 | 5.1 |
| Format C | 5.4 | 86 | 151.9 | 5.6 |
| Format D | 8.0 | 179 | 130 | 5.3 |

**Fig. S1. TcE-characterization.** (**A**) Absolute size exclusion chromatography analysis of TcEs. (**B**) Absolute hydrophobic interaction chromatography analysis of TcEs. (**C**) Differential scanning calorimetry analysis of TcEs. (**D**) Surface plasmon resonance sensograms of TcEs at three different concentrations of CD3ε and (**E**) Her2. Black lines = raw data, red lines = model fitted (**F**) Summary TcE physical characteristics obtained by multi-angle light scattering, together with size exclusion chromatography, surface plasmon resonance and dynamic light scattering. Values in brackets indicate hydrodynamic radii as measured by dynamic light scattering. (**G**) Optimized raw SAXS data curves (colored circles) with error bars (grey lines) and SAXS curves with optimal range (red lines), used for further analysis. (**H**) Examples of multi-state model fits for every TcE or TcE-antigen complex showing best and worst fits. Red lines = experimental SAXS curves with optimal range, blue line = model fit. q($Å^{-1}$) = scattering vector per Angstrom, logI(q) = scattering intensity.

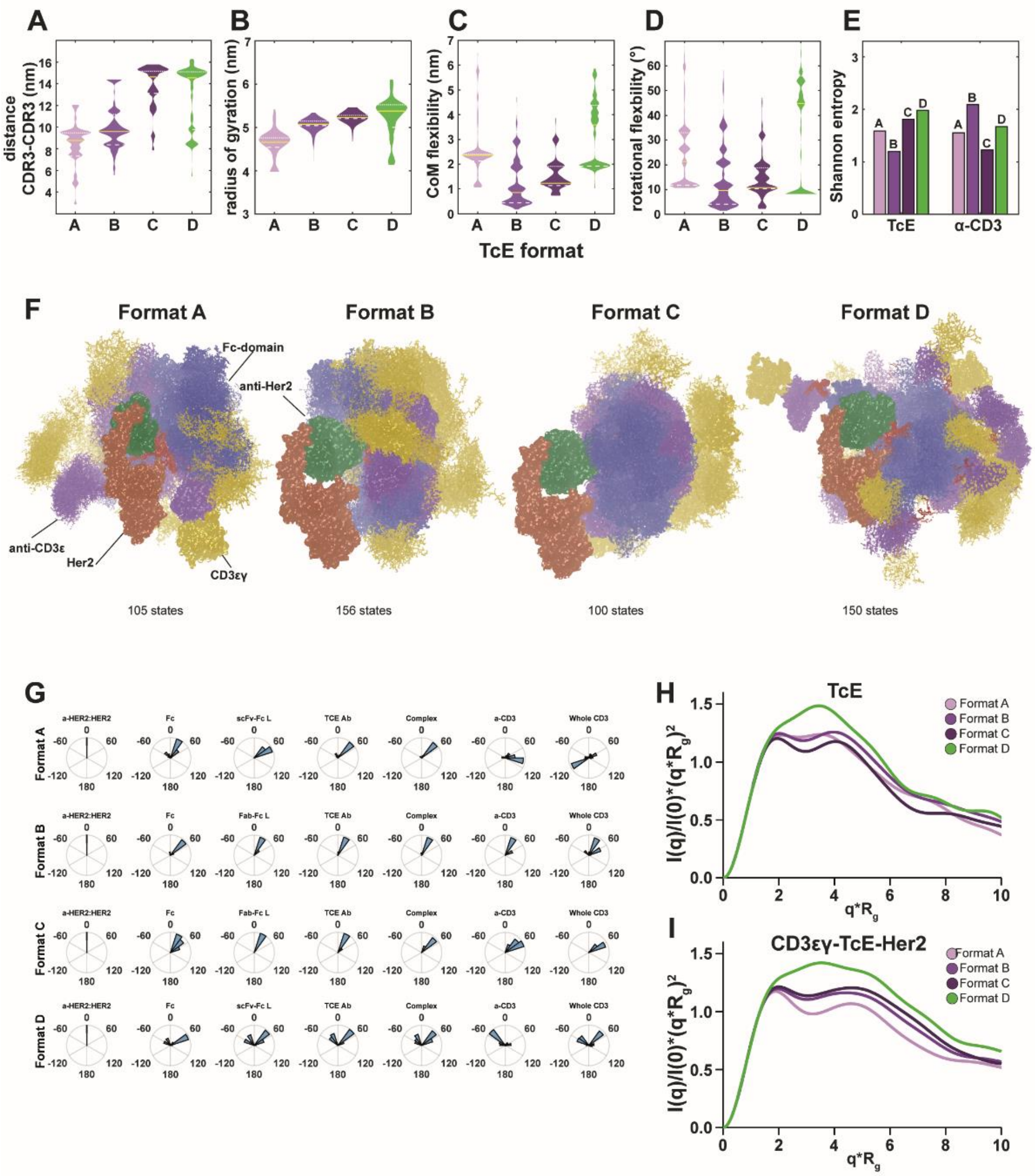

**Fig. S2. Structural and visual characterization of TcE formats.** (**A**) SAXS-data derived conformation weight-adjusted distributions of paratope spacings, (**B**) radius of gyration, (**C**) center of mass (CoM) flexibility. (**D**) rotational flexibility and (**E**) Shannon entropy. Solid yellow lines indicate medians, white dotted lines represent 1st and 3rd quartiles respectively. (**F**) Visual representations of the weight adjusted unique states (PDB structures) in the conformational space for TcE-antigen complexes in solution. The unique PDB models were aligned to anti-Her2 in complex with Her2 and depicted with weighted lines in accordance with their effective weights in the conformational space. Magenta = anti-CD3, green = anti-Her2, blue = Fc-domain, yellow = CD3εγ, salmon = Her2. Flexible scFv-Fc (Format A, Format D) or Fab-Fc linkers (Format B, Format C) are shown in red. (**G**) Conformation weight-adjusted polar histograms depicting tilt angle distributions for each defined domain relative to the Her2–anti-Her2 axis, shown for anti-Her2-Her2 complex, Fc, scFv-Fc/Fab-Fc flexible linkers, TcE Abs embedded within the TcE-antigen complexes, entire TcE-antigen complexes, anti-CD3, and whole CD3. The magnitude of the polar angles represents the angular displacement from the Her2–anti-Her2 axis. The angles were assigned a negative sign if the center of mass of the domain had a negative y-component in the local right-handed coordinate system. (**H**) Dimensionless Kratky plots of isolated TcEs or (**I**) TcE-antigen complexes.

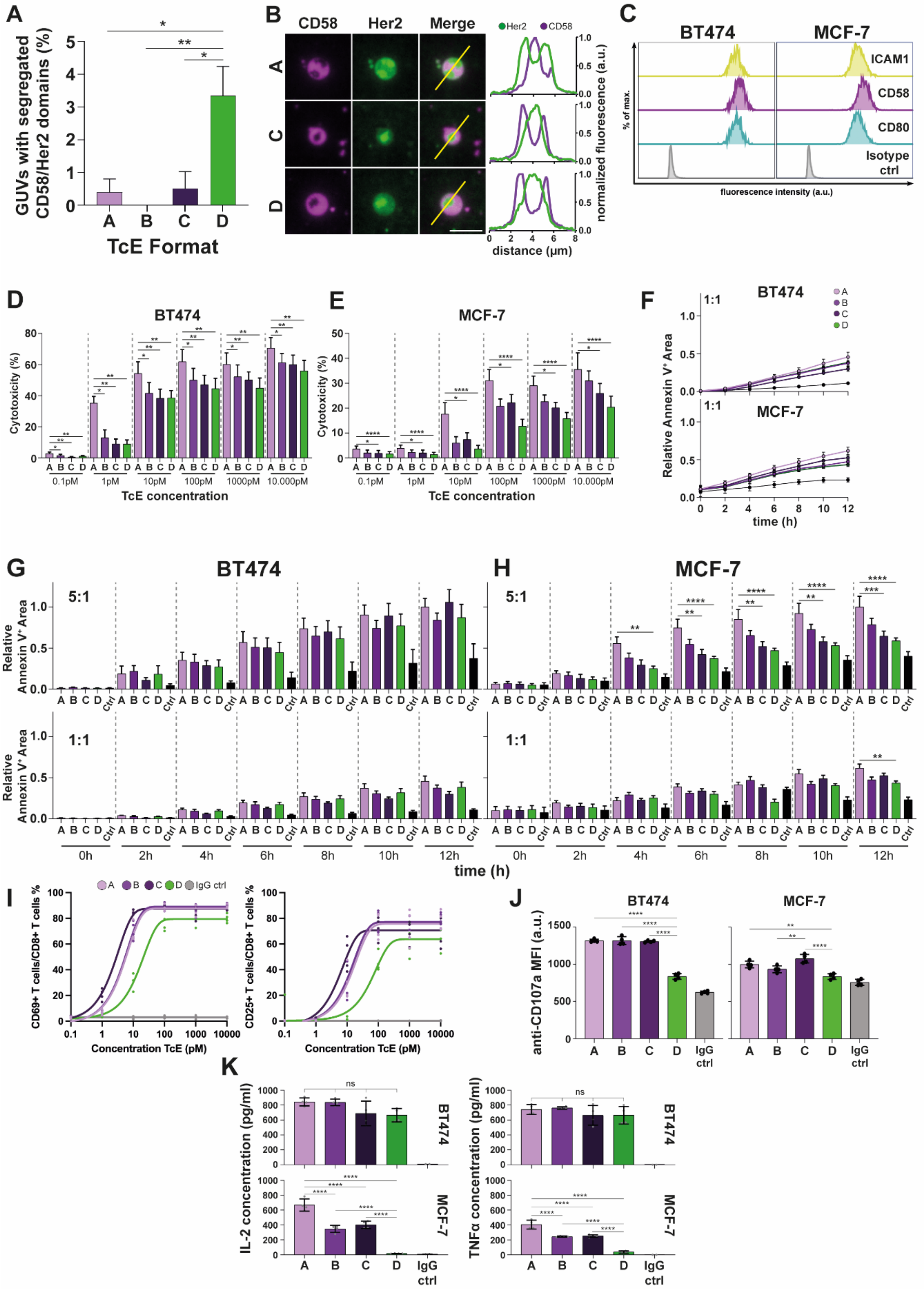

**Fig. S3. CD2-CD58 segregation in GUVs, TcE-mediated cancer cell cytotoxicity, T cell activation and cytokine production.** (**A**) Frequency of GUVs showing segregated domains of CD58 and Her2. Four biological replicates, one-way ANOVA, Mean + SEM, * p≤0.0146, ** p=0.0067. (**B**) Representative TIRF images of CD3 and CD2 loaded GUVs showing segregated domains of CD58 (magenta) and Her2 (green) recruitment. Line plots on the right show normalized fluorescence of CD58 and Her2 along the indicated yellow line, scale bar = 5µm. (**C**) Flow cytometry analysis of BT474 (left) or MCF-7 (right) cancer cell lines stained for ICAM1, CD80, CD58 or isotype control. (**D**) Two-way ANOVA of cytotoxicity of BT474 or MCF-7 (**E**) cancer cells for indicated formats and concentrations corresponding to Fig.2A & B, 6 biological replicates, Median + SEM. * $p<0.0328$, ** $p \leq 0.0012$, **** $p<0.0001$. (**F**) Relative Annexin $5^+$ Area over 12 h in co-cultures of T cells with BT474 (top) or MCF-7 (bottom) cancer cells at a 1:1 E:T ratio in the presence of 50pM TcEs. 3 biological replicates. Solid circles = Median ± SEM. (**G**) Two-way ANOVA of Relative Annexin $5^+$ Area of BT474 or MCF-7 (**H**) cancer cells at a 5:1 (top) or 1:1 (bottom) E:T ratio corresponding to (F) and Fig. 2C & D. (**I**) Flow cytometry analysis of T cell activation markers CD69 (left) and CD25 (right) on T cells after 24h of co-culture with BT474 tumour cells upon increasing TcE concentrations. (**J**) Flow cytometry analysis of surface exposed CD107a on T cells after 21h of co-culture with BT474 (left) or MCF-7 (right) tumor cells in the presence of 5 pM TcEs or IgG control. Mean ± SD. 2 biological replicates. One-way ANOVA, ** $p \leq 0.0044$, *** $p=0.0006$, **** $p<0.0001$. (**K**) IL-2 (left) and TNFα (right) concentrations in the supernatants of T cell-BT474 (top) or MCF-7 (bottom) co-cultures after 24h and in the presence of 5pM TcEs. Mean ± SD. 2 biological replicates. One-way ANOVA, **** $p<0.0001$, ns = not significant.

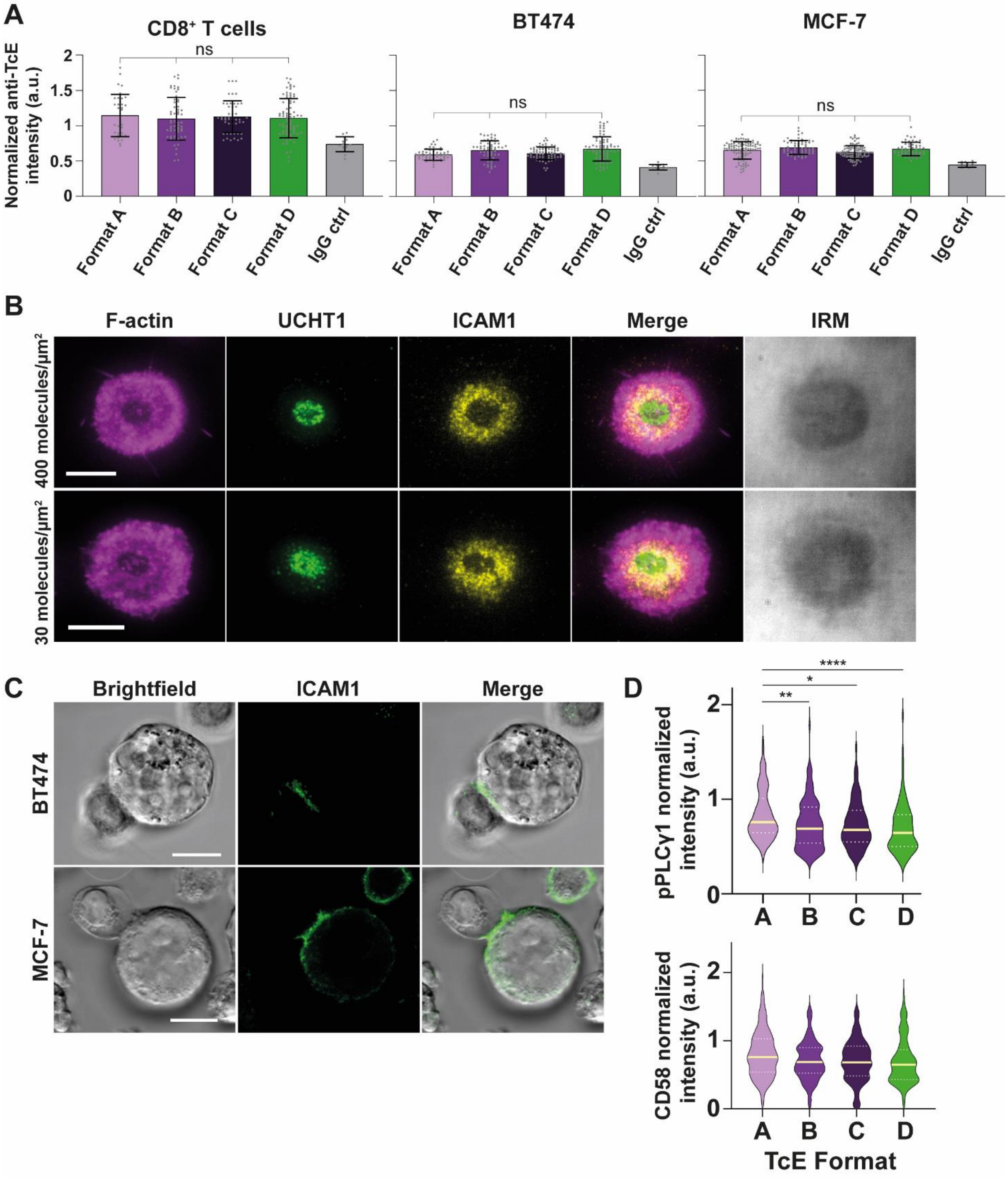


**Fig. S4. TcE-cell binding, UCHT1-mediated IS formation, ICAM1 localization in T cell-tumor cell doublets and PLCγ1 signaling at low Her2 density** (**A**) Quantification of anti-human Fc-PE intensities from confocal microscopy images for TcE formats and isotype control. Mean ± SD, ns = not significant. 2 biological replicates. (**B**) Representative TIRF and IRM images of T cells incubated for 20min. on SLBs presenting 400 (top) or 30 (bottom) molecules/µm$^2$ of UCHT1 (green) and ICAM1 (yellow) at 200 molecules/µm$^2$, scale bars = 8µm. Note that UCHT1 is shown with different brightness settings to avoid saturation (**C**) Representative single-plane confocal and brightfield images of T cell-BT474 (top) or MCF-7 (bottom) duplets after 2h of co-incubation in the presence of 5pM Format A and staining for ICAM1, scale bars = 10µm. (**D**) Normalized pPLCγ1 (top) and CD58 (bottom) levels at the TcE-mediated IS at 30 (Her2) and 200 (ICAM1) molecules/µm$^2$ in the presence of 200 molecules/µm$^2$ CD58. Yellow solid line = median, White dashed line = quartiles. 2 biological replicates, Kruskal-Wallis test, * $p<0.05$, ** $p<0.01$, **** $p<0.0001$.

**Movies S1-4 (separate files).** Visual representations of the weight adjusted unique states (PDB structures) in the conformational space for Format A (S1), Format B (S2), Format C (S3), Format D (S4) while in complex with CD3 and Her2 in solution. The unique PDB models were aligned to anti-Her2 in complex with Her2 and depicted with weighted lines in accordance with their effective weights in the conformational space. Magenta = anti-CD3, green = anti-Her2, blue = Fc-domain, yellow = CD3εγ, salmon = Her2. Flexible scFv-Fc (Format A, Format D) or Fab-Fc linkers (Format B, Format C) are shown in red.

**Movie S5 (separate file).** BT474 cancer cell (CFSE labelled, magenta) and $CD8^+$ T cell co-culture in an E:T ratio of 5:1 over a time-course of 12h in the presence or absence of 50 pM TcEs and fluorescently labelled Annexin V (cyan). Scale bar = 500µm.

**Movie S6 (separate file).** MCF-7 cancer cell (CFSE labelled, magenta) and $CD8^+$ T cell co-culture in an E:T ratio of 5:1 over a time-course of 12h in the presence or absence of 50 pM TcEs and fluorescently labelled Annexin V (cyan). Scale bar = 500µm.
